# Direct Observation of Stepping Rotation of V-ATPase Reveals Rigid Coupling between V_o_ and V_1_ Motors

**DOI:** 10.1101/2022.06.13.494302

**Authors:** Akihiro Otomo, Tatsuya Iida, Yasuko Okuni, Hiroshi Ueno, Takeshi Murata, Ryota Iino

**Affiliations:** Institute for Molecular Science, National Institutes of Natural Sciences, 5-1 Higashiyama, Myodaiji-Cho, Okazaki, Aichi 444-8787, Japan; Department of Functional Molecular Science, School of Physical Sciences, SOKENDAI (Graduate University for Advanced Studies), Shonan Village, Hayama, Kanagawa 240-0193, Japan; Department of Applied Chemistry, Graduate School of Engineering, The University of Tokyo, 7-3-1 Hongo, Bunkyo-Ku, Tokyo 113-8656, Japan; Department of Chemistry, Graduate School of Science, Chiba University, 1-33 Yayoi-Cho, Inage-Ku, Chiba 263-8522, Japan

**Author notes:** Ryota Iino **Email:**. **Author Contributions:** H.U., T.M. and R.I. conceived original concept; A.O. and R.I. designed research; A.O. performed research and data analysis; A.O., T.I., Y.O., H.U., and T.M. contributed to sample preparation; and A.O. and R.I. wrote the paper. **Competing Interest Statement:** The authors declare no competing interest.

**Keywords:** V-ATPase, Single-molecule analysis, Molecular motors

## Abstract

V-ATPases are rotary motor proteins which convert chemical energy of ATP into electrochemical potential of ions across the cell membrane. V-ATPases consist of two rotary motors, V_o_ and V_1_, and *Enterococcus hirae* V-ATPase (EhV_o_V_1_) actively transports Na^+^ in V_o_ (EhV_o_) by using torque generated by ATP hydrolysis in V_1_ (EhV_1_). Here, we observed ATP-driven stepping rotation of detergent-solubilized EhV_o_V_1_ wild-type, aE634A, and BR350K mutants under the various Na^+^ and ATP concentrations ([Na^+^] and [ATP], respectively) by using a 40-nm gold nanoparticle as a low-load probe. When [Na^+^] was low and [ATP] was high, under the condition that only Na^+^ binding to EhV_o_ is the rate-limiting, wild-type and aE634A exhibited 10-pausing positions reflecting 10-fold symmetry of the EhV_o_ rotor and almost no backward steps. Duration time before forward steps was inversely proportional to [Na^+^], confirming that Na^+^ binding triggers the steps. When both [ATP] and [Na^+^] were low, under the condition that both Na^+^ and ATP bindings are rate-limiting, aE634A exhibited 13-pausing positions reflecting 10- and 3-fold symmetries of EhV_o_ and EhV_1_, respectively. Distribution of duration time before forward step was well fitted by a sum of two exponential decay functions with distinct time constants. Furthermore, frequent backward steps smaller than 36° were observed. Small backward steps were also observed during long, three ATP cleavage pauses of BR350K. These results indicate that EhV_o_ and EhV_1_ do not share pausing positions and Na^+^ and ATP bindings occur at different angles, and the coupling between EhV_o_ and EhV_1_ is not elastic but rigid.

**Significance Statement:** V-ATPases are ion pumps consisting of two rotary motor proteins V_o_ and V_1_, and actively transport ions across the cell membrane by using chemical energy of ATP. To understand how V-ATPases transduce the energy in the presence of structural symmetry mismatch between V_o_ and V_1_, we simultaneously visualized rotational pauses and forward and backward steps of V_o_ and V_1_ coupled with ion transport and ATP hydrolysis reaction, respectively. Our results indicate rigid coupling of a V-ATPase which has multiple peripheral stalks, in contrast to elastic coupling of F-ATPases with only one peripheral stalk, which work as ATP synthase. Our high-speed/high-precision single-molecule imaging of rotary ATPases in action will pave the way for a comprehensive understanding of their energy transduction mechanisms.

## Introduction

Rotary ATPases are ubiquitously expressed in living organisms and play important roles in biological energy conversions (1-6). These rotary ATPases are classified into F-, V-, and A-ATPases based on their amino acid sequences and physiological functions (6). Eukaryotic and bacterial F-ATPases (F_o_F_1_), and archaeal A-ATPases (A_o_A_1_) mainly function as ATP synthases driven by electrochemical potential of ion across the cell membrane, although they can also act as active ion pumps driven by ATP hydrolysis depending on the cellular environments. On the other hand, V-ATPases (V_o_V_1_) in eukaryotes primarily function as active ion pumps. V-ATPases are also found in bacteria, and some of them are termed as V/A-ATPases based on their physiological function, ATP synthesis (6, 7).

To date, numerous studies have been conducted to understand how the two motor proteins (i.e., F_1_/A_1_/V_1_ and F_o_/A_o_/V_o_) of the rotary ATPases couple their rotational motions and functions. Single-molecule studies using fluorescent probes (8-11), gold nanoparticle (AuNP) or nanorod probes (12-20), and Förster resonance energy transfer (FRET) (15, 21, 22) have revealed rotational dynamics of the rotary ATPases for both ATP hydrolysis/synthesis directions. Furthermore, recent cryo-electron microscopic (cryo-EM) single-particle analyses revealed entire architectures of the rotary ATPases with different structural states at atomic resolutions (23-34). In particular, several studies have demonstrated elastic coupling of F_o_F_1_ due to large deformations of the peripheral stalk connecting F_o_ and F_1_ (24, 28, 34). However, few studies on other types of rotary ATPases with different functions and subunit compositions have been performed, and a comprehensive understanding of the energy transduction mechanism remains elusive.

*Enterococcus hirae* V-ATPase (EhV_o_V_1_) works as an ATP-driven sodium ion (Na^+^) pump to maintain Na^+^ concentration ([Na^+^]) inside the cell (Figure 1A) (35-39). Note that we use the term V-ATPase or V_o_V_1_ because its physiological function is not ATP synthesis but active ion transport. EhV_o_V_1_ is a multi-subunit complex composed of 9 different subunits, namely, ac_10_dE_2_G_2_ and A_3_B_3_DF complexes in EhV_o_ and EhV_1_, respectively. In EhV_1_ A_3_B_3_DF complex, three pairs of the A and B subunits form a hetero-hexameric A_3_B_3_ stator ring, and the central rotor DF subcomplex is inserted into the A_3_B_3_ ring (Figure 1B, bottom) (40, 41). The EhV_o_ ac_10_dE_2_G_2_ complex transports Na^+^ across the cell membrane. The membrane-embedded rotor ring is formed by a decamer of tetra-helical transmembrane c-subunit (c_10_-ring, Figure 1B, top), connected with the central DF stalk via the d-subunit (25, 42). The stator a-subunit works as an ion channel, and two EG peripheral stalks interact with the a-subunit and A_3_B_3_ ring to assure rotary coupling between EhV_o_ and EhV_1_. In EhV_1_, the ATP hydrolysis reaction is catalyzed at the interfaces of three A and B subunits. It drives a counterclockwise rotation of the DF rotor subunits as viewed from EhV_o_ side (Figure 1B, bottom). Likewise other F_1_/A_1_/V_1_ (10, 43), EhV_1_ is a stepping motor that rotates 120° per one ATP hydrolysis (44, 45). By using high-speed and high-precision single-molecule imaging analysis with AuNP as a low-load probe, we previously revealed that the 120° step of isolated EhV_1_ is further divided into 40 and 80° substeps, which are triggered by ATP binding and ADP release, respectively (44). In contrast, as our previous single-molecule observation of EhV_o_V_1_ did not resolve clear pauses and steps (16), elementary steps in the rotation of EhV_o_V_1_ have not been revealed.

**Figure 1.**
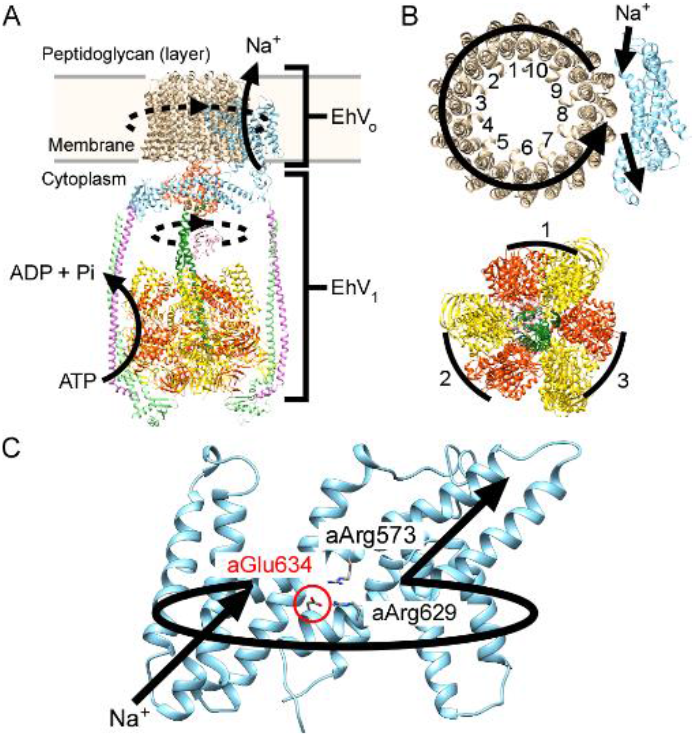
(*A*) Overall architecture of EhV_o_V_1_. The dotted circular arcs represent the rotation direction driven by ATP hydrolysis. (*B*) (top) Top view of a-subunit (cyan) and c_10_-ring (brown) of EhV_o_ and (bottom) A-(yellow), B-(orange), D-(green), and F-subunits (pink) of EhV_1_. The black arrow at the top indicates the path of Na^+^ movement during ATP-driven rotation. The arcs at the bottom represent the catalytic AB pairs. (*C*) Side view of a-subunit viewed from c-subunit. This structure was constructed by a SWISS-MODEL server (80) by using a structure of a-subunit of V-ATPase from *Thermus Thermophilus*. The black arrows represent the path of Na^+^ movement during ATP-driven rotation. The mutated residue, aGlu634, is located on the surface of the entry half-channel of a-subunit as highlighted in red letters and a circle.

Although the mechanism of ion transport in the F_o_/A_o_/V_o_ has not been fully understood, the so-called “two-channel” model has been widely accepted (46-52). In this model, the a-subunit has two half-channels for ion entry/exit into/from the ion-binding sites of rotor c-ring. In the case of EhV_o_, Na^+^ enters the half-channel from the cytoplasmic side and binds to the negatively charged Na^+^-binding sites of the c-subunit (42, 53). Then, the charge-neutralized c-subunit can move into the hydrophobic lipid membrane (51, 54). The rotational torque generated by ATP hydrolysis in EhV_1_ is transmitted to EhV_o_ via the rotor d-subunit, allowing the c_10_-ring to rotate unidirectionally in the lipid membrane. Na^+^ translocated by a nearly single turn of the c_10_-ring reaches another half-channel of the a-subunit, which connects the Na^+^-binding site of c-subunit to the extracellular side. Then, Na^+^ is pumped out of the cell by a hydrated microenvironment (55) and/or the electrostatic repulsion with the positively charged residues in the a-subunit, aArg573 and aArg629, located at the interface between two half-channels (Figures 1C and S1) (25). Because EhV_o_V_1_ has the c_10_-ring, ten Na^+^ are transported per single turn. Therefore, the step size of EhV_o_ is expected to be 36° (360°/10), similar to *Escherichia coli* (*E. coli*) and yeast F_o_F_1_, which also have c_10_-rings (12, 13, 21, 34).

The ion-to-ATP ratio is a central issue in the coupling mechanism of rotary ATPases. All known F_1_/A_1_/V_1_ has three catalytic sites and 3-fold structural symmetry, and hydrolyzes or synthesizes 3 ATP molecules per single turn. In contrast, the number of protomers forming the rotor c-ring of F_o_/A_o_/V_o_ varies from 8 to 17 depending on the species, suggesting wide variations in the ion-to-ATP ratio of rotary ATPases (56, 57). In EhV_o_V_1_, because the rotor c-ring of EhV_o_ has a 10-fold structural symmetry (Figure 1B, top), this enzyme has a structural symmetry mismatch and a non-integer ratio between transported Na^+^ and hydrolyzed ATP (10/3 = 3.3). If the rotational coupling between EhV_o_ and EhV_1_ is elastic, as reported for *E. coli* and yeast F_o_F_1_, the symmetry mismatch is relieved by large deformations of the peripheral stalk and/or the central rotor (24, 28, 34, 58). On the other hand, if the coupling is rigid due to multiple peripheral stalks of the EhV_o_V_1_, the pausing positions of both EhV_o_ and EhV_1_ would be observed independently in a single-molecule observation. To address this issue, it is required to directly visualize the rotational pauses and steps of EhV_o_V_1_ under the conditions that elementary steps of rotation for EhV_o_ and EhV_1_ are both rate-limiting, for example, the bindings of Na^+^ and ATP to EhV_o_ and EhV_1_, respectively.

Here we carried out high-speed and high-precision single-molecule imaging of rotation of detergent-solubilized EhV_o_V_1_ by using 40-nm AuNP as a low-load probe. To resolve the rotational pauses and steps of EhV_o_, a glutamate residue in the stator a-subunit (aGlu634) was replaced with alanine. Since the mutated aGlu634 is located on the surface of Na^+^ entry half-channel (Figures 1C and S2), Na^+^ binding to the c-subunit in EhV_o_V_1_(aE634A) mutant (hereinafter, referred to as aE634A) is expected to become slower than in wild-type. The rotation rate of aE634A decreased about 10 times compared with that of the wild-type, allowing us to clearly resolve the rotational pauses and steps in EhV_o_. Under the condition that only Na^+^ binding is the rate-limiting, aE634A showed 10 pausing positions per single turn and a step size of about 36°, consistent with 10 protomers in the c_10_-ring of EhV_o_. The duration time before the forward step was inversely proportional to the [Na^+^], indicating that the dwell corresponds to the waiting time for Na^+^ binding. On the other hand, under the condition that both Na^+^ and ATP bindings are the rate-limiting, 13 pausing positions per single turn were observed. Furthermore, backward steps smaller than 36° were frequently observed only when ATP binding is also the rate-limiting, indicating that EhV_o_V_1_ undergoes Brownian motion between adjacent pausing positions of EhV_o_ and EhV_1_ when no torque is applied from EhV_1_. The backward steps of 36° or larger than 36° were rarely observed, suggesting the suppression of the reverse Na^+^ transport. Small backward steps were also frequently observed during long, three ATP cleavage pauses of another mutant EhV_o_V_1_(BR350K) in which ATP hydrolysis is the rate-limiting of rotation (44). From these results, we conclude that EhV_o_ and EhV_1_ do not share their pausing positions and Na^+^ and ATP bindings occur at different angles, and their coupling is not elastic but rigid.

## Results

### Single-molecule imaging system and generation of EhV_o_V_1_ aE634A mutant

For the single-molecule observation, a poly-histidine tag (His_3_-tag) was introduced to the C-terminus of c-subunit to immobilize the detergent-solubilized EhV_o_V_1_ complex on a Ni^2+^-NTA coated glass surface (Figure 2A). Furthermore, the N-terminus of A-subunit was biotinylated by a genetically fused Avi-tag (59), and streptavidin-coated AuNP probe (*φ*=40 nm) was attached (Figure 2A). Note that the stator subunits rotate against the rotor subunits by ATP hydrolysis in our experimental system because the rotor c_10_-ring was fixed on the glass surface. The ATP-driven rotation was observed under the total internal reflection dark-field microscope at 1,000 or 3,000 frames per second (fps) at 25°C. In this system, the localization precision was 0.6 nm at 3,000 fps (0.33 ms temporal resolution), determined by centroid analysis of the scattering images of single AuNPs non-specifically attached to the glass surface (Figure S3).

**Figure 2.**
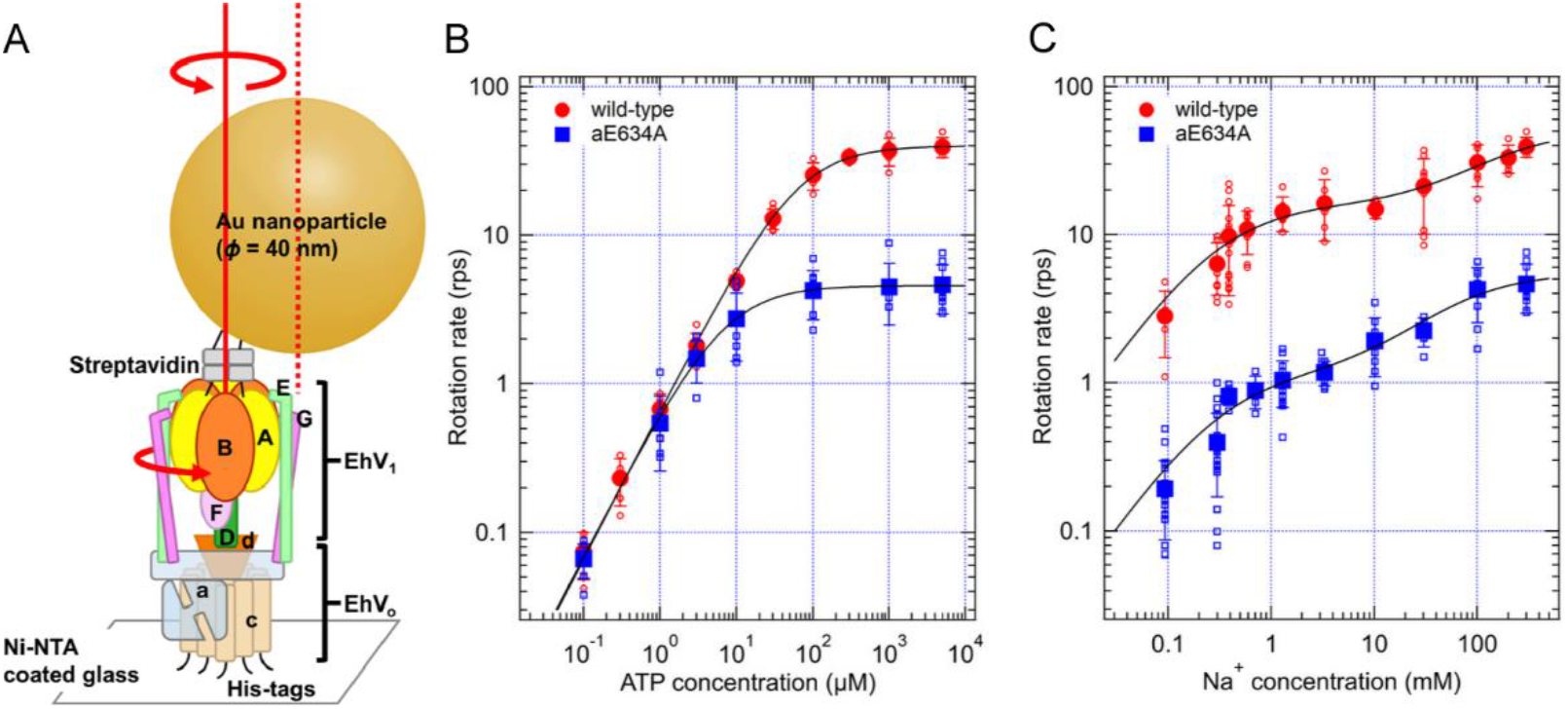
(*A*) Schematic model of single-molecule rotation assay of EhV_o_V_1_ probed with AuNP. Each alphabet represents the name of subunits. EhV_o_V_1_ was fixed on the Ni^2+^-NTA-coated cover glass via His_3_-tags added to the C-terminus of c-subunit. The streptavidin-coated gold nanoparticle (40 nm in diameter) was attached to the N-terminus of A-subunit which was biotinylated by adding Avi-tag. Solid and dotted red lines indicate the center axis of EhV_o_V_1_ and the centroid of attached AuNP, respectively. Because the rotor c_10_-ring was fixed on a glass surface, the stator subunits rotate counterclockwise against rotor subunits as shown by red arrows. (*B*) [ATP] dependence of rotation rate of wild-type and aE634A at 300 mM NaCl. Red open and blue open squares indicate the data from individual molecules of wild-type and aE634A, respectively. The closed symbols are average, and error bar is the standard deviation. Data were fitted with Michaelis-Menten equation: *V*=*V*_max_^ATP^·[ATP]/(*K*_m_^ATP^+[ATP]). The obtained kinetic parameters are summarized in Table 1. (*C*) [Na^+^] dependence of rotation rate of wild-type and aE634A at 5 mM ATP. The correspondence of colored symbols is the same as in *B*. The black lines show the fit with sum of two-independent Michaelis-Menten equations: *V*=*V*_max1_^Na^·[Na^+^]/(*K*_m1_^Na^+[Na^+^]) + *V*_max2_^Na^·[Na^+^]/(*K*_m2_^Na^+[Na^+^]). The obtained kinetic parameters are summarized in Table 2. The contaminated Na^+^ in the observation buffers were taken into account as shown in Figure S6. Namely, 50 mM Bis-Tris (pH6.5) was used for 0.09 mM Na^+^, and 20 mM potassium phosphate (pH6.5) was used for other [Na^+^]s as buffers.

To clearly visualize the ATP-driven rotation rate-limited by Na^+^ transport, site-directed mutagenesis was conducted on the a-subunit of EhV_o_. In the a-subunit of V_o_, there are some negatively charged amino acid residues on the surface of half-channels forming ion transport pathways (23, 60). In EhV_o_, aGlu634 is located on the Na^+^ entry half-channel (Figure 1C). This glutamate residue is completely conserved among other V_o_V_1_ (Figure S1), suggesting that this negatively charged residue has a role in attracting ions into the entry half-channel. To reduce the Na^+^ binding rate, we prepared EhV_o_V_1_ mutant, aE634A, where aGlu634 was replaced with alanine to eliminate the negative charge of the side chain. The purified aE634A solubilized with *n*-dodecyl-β-D-maltoside (DDM) exhibited the subunit stoichiometry similar to the wild-type in SDS-PAGE (Figure S4), indicating intactness of the complex.

### [ATP] and [Na^+^] dependence of rotation rate of wild-type and aE634A

Rotation rates of wild-type and aE634A at various substrate concentrations were examined from the slope of the time course of rotation (Figures 2B, C, and S5). [ATP] dependences of the rotation rate for wild-type and aE634A in the presence of 300 mM Na^+^ are shown in Figure 2B. Under this high [Na^+^] condition, Na^+^ binding is fast and not rate-limiting, and consequently, [ATP] dependence obeyed Michaelis-Menten kinetics. Obtained kinetic parameters, the Michaelis constant (*K*_m_^ATP^) and the maximum velocity (*V*_max_^ATP^), are summarized in Table 1. The value of *K*_m_^ATP^ for wild-type was 60.4 ± 2.1 μM (fitted value ± S.E. of the fit). This value was comparable to that for isolated EhV_1_ (43 ± 6 μM), indicating that apparent affinity of ATP was not significantly affected by EhV_o_. On the other hand, the value of *V*_max_^ATP^ (40.0 ± 0.3 rps, fitted value ± S.E. of the fit) was significantly smaller than that of the isolated EhV_1_ (117 ± 3 rps) (44). As a result, the binding rate constant of ATP (*k*_on_^ATP^) estimated by 3 × *V*_max_^ATP^ / *K*_m_^ATP^ was smaller than that of the isolated EhV_1_ (Table 1). We attributed this difference to the intact interaction between rotor c_10_-ring and stator a-subunit of EhV_o_ and concluded that Na^+^ transport in EhV_o_ limits the *V*_max_^ATP^ of EhV_o_V_1_, as discussed in our previous single-molecule study (16). In aE634A, the values of *K*_m_^ATP^ and *V*_max_^ATP^ were 6.6 ± 0.3 μM and 4.58 ± 0.04 rps, respectively, and both were approximately one-tenth of those for wild-type (Table 1). Thus, the *k*_on_^ATP^ for aE634A (2.1 × 10^6^ M^-1^s^-1^) was in close agreement with that for wild-type (2.0 × 10^6^ M^-1^s^-1^) (Table 1). These results indicate that the mutation in aGlu634 affects only Na^+^ transport but not ATP binding in the EhV_o_V_1_ complex.

**Table 1.**
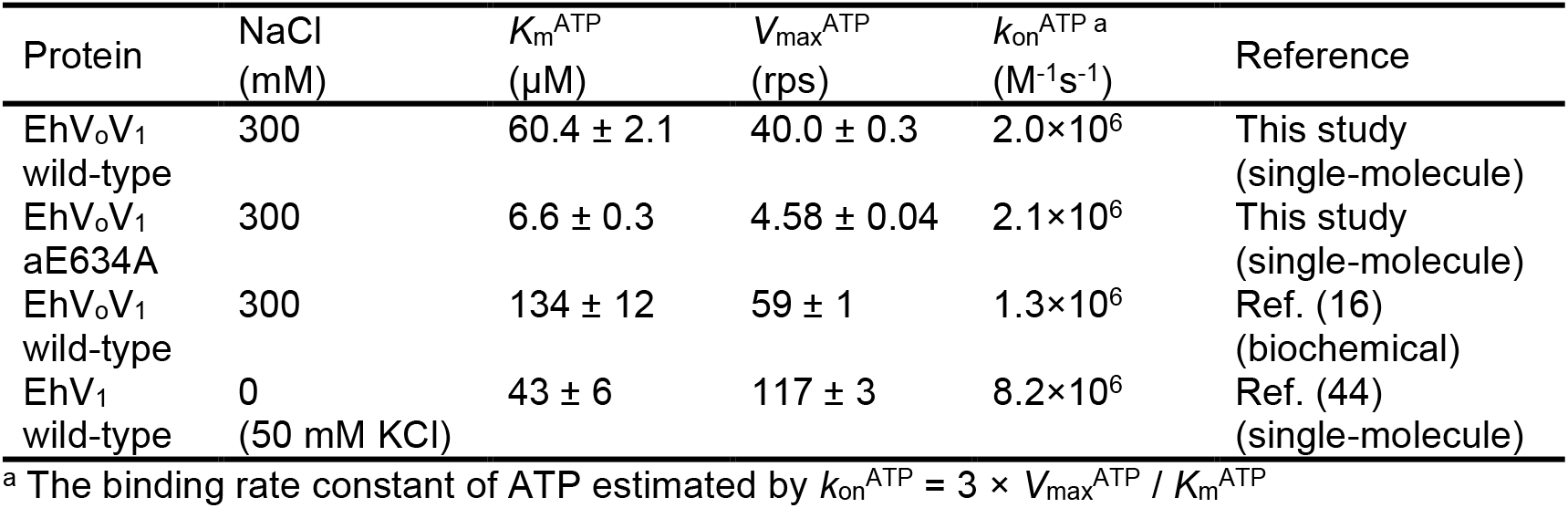
Kinetic parameters for [ATP] dependence of rotation rate of EhV_o_V_1_ and EhV_1_

Figure 2C shows [Na^+^] dependence of the rotation rate at the saturated [ATP] (5 mM). Here, contaminated Na^+^ in the observation buffers were estimated by an inductivity coupled plasma optical emission spectrometer (ICP-OES) (Figure S6), and total [Na^+^] including contamination was used for the plot. Under this condition, [ATP] is sufficiently high and ATP binding is not rate-limiting (Figure 2B and Table 1), and the effect of [Na^+^] on the rotation rate becomes obvious. As a result, both wild-type and aE634A showed biphasic responses to [Na^+^]. As reported in our previous studies, data were fitted by a summation of two independent Michaelis-Menten equations assuming two Na^+^ binding sites (16, 61). The obtained kinetic parameters are shown in Table 2. The both values of *V*_max1_^Na^ and *V*_max2_^Na^ for aE634A (1.1 ± 0.3 and 4.4 ± 0.5 rps, fitted value ± S.E. of the fit) were about ten times smaller than those for wild-type (15.7 ± 1.6 and 35.0 ± 7.1 rps) (Table 2). On the other hand, considering the large fitting error especially in wild-type, the values of *K*_m1_^Na^ and *K*_m2_^Na^ do not seem to be significantly different between aE634A (0.32 ± 0.28 and 58.3 ± 26.5 mM, fitted value ± S.E. of the fit) and wild-type (0.29 ± 0.09 and 160.2 ± 88.8 mM). Then, the binding rate constant of Na^+^ (*k*_on_^Na^) estimated by 10 × *V*_max_^Na^ / *K*_m_^Na^ largely decreased in aE634A, especially more than ten times for the high-affinity site (*k*_on1_^Na^, Table 2). The significant decrease in the *V*_max2_^Na^ for aE634A compared with that of wild-type also suggests that the rate of elementary reaction steps other than Na^+^ binding also decreased in EhV_o_. Because Na^+^ binding to EhV_o_ is certainly the rate-limiting at low [Na^+^] for both wild-type and aE634A, we then conducted a detailed analysis of the rotational pauses and steps at low [Na^+^].

**Table 2.** Kinetic parameters for [Na^+^] dependence of rotation rate of wild-type and aE634A ^a^

| Protein | $K_{m1}^{\text{Na}}$<br>(mM) | $V_{\text{max}1}^{\text{Na}}$<br>(rps) | $k_{\text{on}1}^{\text{Na} \text{ b}}$<br>(M<sup>-1</sup>s<sup>-1</sup>) | $K_{m2}^{\text{Na}}$<br>(mM) | $V_{\text{max}2}^{\text{Na}}$<br>(rps) | $k_{\text{on}2}^{\text{Na} \text{ b}}$<br>(M<sup>-1</sup>s<sup>-1</sup>) |
| --- | --- | --- | --- | --- | --- | --- |
| EhV<sub>o</sub>V<sub>1</sub><br>wild-type | 0.29 ± 0.09 | 15.7 ± 1.6 | 5.4 × 10<sup>5</sup> | 160.2 ± 88.8 | 35.0 ± 7.1 | 2.2 × 10<sup>3</sup> |
| EhV<sub>o</sub>V<sub>1</sub><br>aE634A | 0.32 ± 0.28 | 1.1 ± 0.3 | 3.4 × 10<sup>4</sup> | 58.3 ± 26.5 | 4.4 ± 0.5 | 7.5 × 10<sup>2</sup> |

<sup>a</sup> [ATP] was 5 mM<sup>b</sup> The binding rate constant of Na<sup>+</sup> estimated by $k_{\text{on}1(2)}^{\text{Na}} = 10 \times V_{\text{max}1(2)}^{\text{Na}} / K_{m1(2)}^{\text{Na}}$

### Pauses and steps of EhV_o_ and EhV_1_ in ATP-driven rotation of wild-type and aE634A

Figures 3A and S7 show typical time courses of rotational trajectory (pink line) of wild-type observed at 0.3 mM Na^+^ and 5 mM ATP with 3,000 fps (0.33 ms temporal resolution). Enlarged trajectory for single revolution and corresponding *x-y* trajectory (inset, pink line) are also shown. Under this condition, not ATP binding but Na^+^ binding is the rate-limiting of ATP-driven rotation (Figures 2B and C, Table 2). The rotational trajectory showed transient pauses and steps in a forward, counterclockwise direction. Then, the pauses and steps were objectively detected with a step-finding algorithm (62) (black line on the trajectory) applied to median-filtered traces (current ± 4 frames, red line), as our previous studies on other motor proteins (63, 64). As indicated by the numbers in the trajectories, 10 pauses were mainly detected in a single turn (Figure S8A), reflecting the number of protomers in rotor c_10_-ring (Figure 1B, top). In addition, number of pauses smaller or larger than 10 per single turn was also detected (Figure S8A). The number of pauses smaller than 10 can be attributed to undetectable short pauses due to the limited time resolution, because stepping behavior is a stochastic process. The number of pauses larger than 10 is presumably attributed to occasional detection of the sub-pauses of EhV_1_ for ADP release, which has a time constant of 2.5 ms and is independent of [ATP] (44). Furthermore, small fraction of under-or over-fitting of the step-finding algorithm would also be included (64). Distribution of the step size revealed that backward steps were rarely observed (0.8%, Figure 3B), the average value of forward step was 36.6°, and a Gaussian function well fitted the distribution with the peak at 33.8 ± 14.2° (peak ± S.D.). These values show excellent agreement with the step size expected from the structural symmetry of c_10_-ring (360°/10=36°). Distribution of the duration time before forward step was well fitted by a single exponential decay function with the time constant of 13.5 ± 0.3 ms (fitted value ± S.E. of the fit), indicating single rate-limiting step of the rotation (Figure 3C).

**Figure 3.**
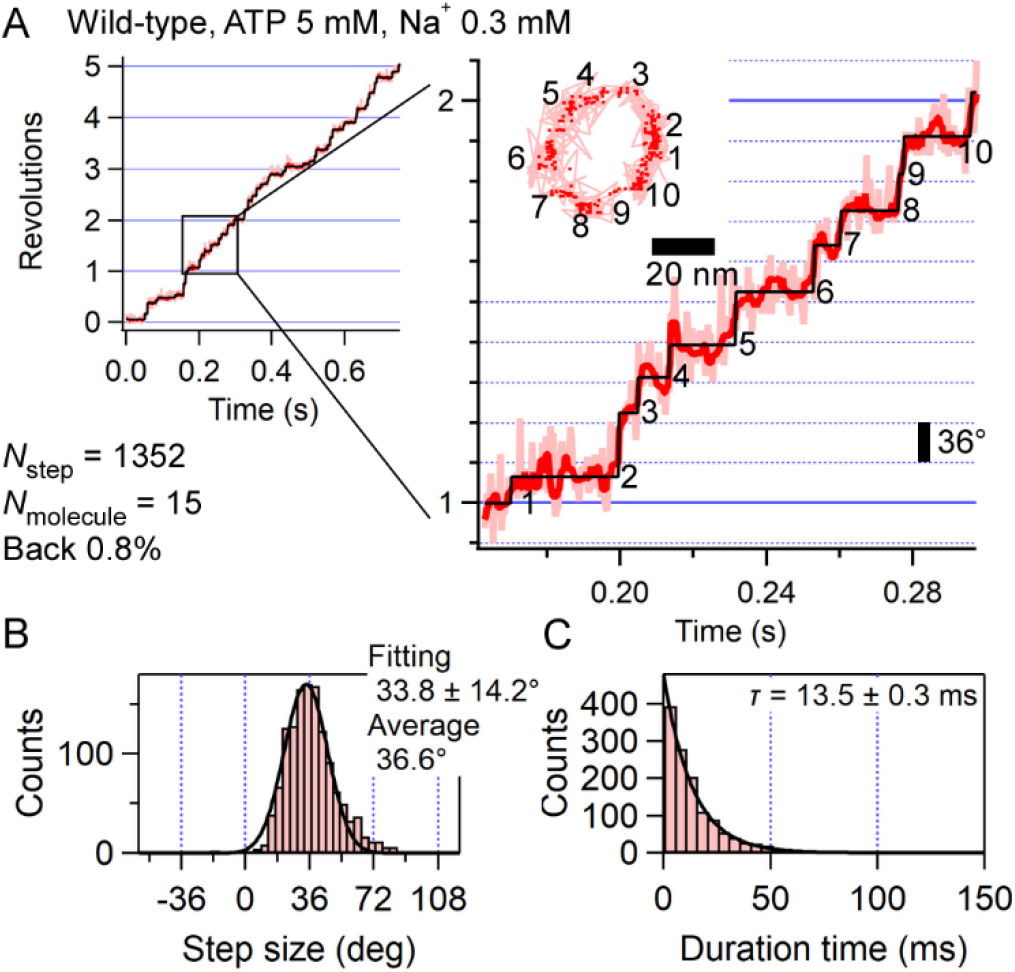
(*A*) Typical trajectory of ATP-driven rotation of wild-type EhV_o_V_1_ at 5 mM ATP and 0.3 mM Na^+^ recorded with 3,000 fps (0.33 ms time resolution). Enlarged view of one revolution (360°) is shown on the right. Pink, red, and black traces represent raw, median-filtered (current ± 4 frames), and fitted trajectories, respectively. Inset shows the corresponding *x-y* trajectory. Pink lines and red dots represent the raw and median-filtered (current ± 4 frames) coordinates, respectively. We collected 1352 steps from 15 molecules. Other examples of trajectory are shown in Figure S7. (*B*) Distribution of the step size fitted with single Gaussian assuming single peak. The right top values are fitted parameter (peak ± S.D.) and average. The ratio of backward steps was 0.8%. (*C*) Distribution of the duration time before forward step fitted with a single exponential decay function. The right top values are obtained time constant (fitted value ± S.E. of the fit).

Rotational trajectories of aE634A observed at 0.3 mM Na^+^ and 5 mM ATP with 1,000 fps are shown in Figures 4A and S9. In aE634A, the median filtering of current ± 7 frames was applied for the step-finding algorithm (62). As in wild-type, the number of pauses per single turn was mainly 10 (Figure S8B), and the step size was 37.1° (average) or 34.9 ± 12.8° (peak ± S.D.) (Figure 4B), consistent with the structural symmetry of c_10_-ring, and the ratio of backward step was minor (1.2%). Distribution of the duration time before forward steps for aE634A was well fitted by a single exponential decay function with the time constant of 137 ± 6 ms (Figure 4C) which is 10 times larger than that for wild-type, consistent with the analysis of rotation rate (Figure 2C and Table 2). For aE634A, we also examined [Na^+^] dependence, and Figure S10 shows rotational trajectories at 0.09 or 1.3 mM Na^+^ and 5 mM ATP. Distributions of the step size and the duration time before forward step at 0.09 and 1.3 mM Na^+^ are shown in Figure 5A-D, respectively. As same at 0.3 mM Na^+^, the step sizes of about 36° were obtained, indicating that 36° step is independent of [Na^+^]. In contrast, although distributions of the duration time before forward step were also well fitted by single exponential decay functions as 0.3 mM Na^+^, the time constants changed largely. Figure 5E shows [Na^+^] dependence of the time constant. The slope was −1.1, in the case that the data point at 1.3 mM Na^+^ was excluded because it is the first saturating concentration observed in the rotation rate analysis (Figure 2C and Table 2). If this data point was included, the slope was −1.0. Therefore, the time constant is almost inversely proportional to [Na^+^], indicating that Na^+^ binding is the rate-limiting of rotation. Furthermore, this result indicates that ATP hydrolysis in EhV_1_ is tightly coupled with Na^+^ transport in EhV_o_ in the EhV_o_V_1_ complex solubilized with DDM.

**Figure 4.**
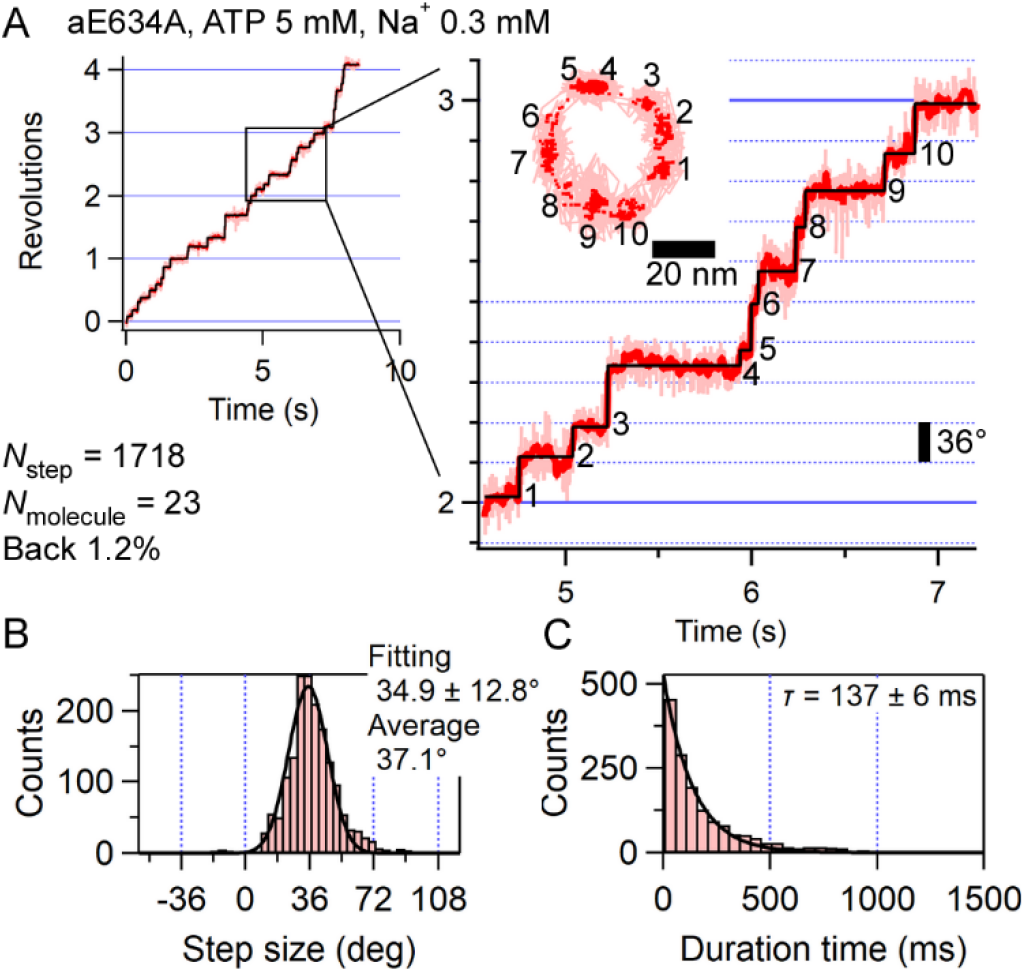
(*A*) Typical trajectory of ATP-driven rotation of aE634A at 5 mM ATP and 0.3 mM Na^+^ recorded with 1,000 fps (1 ms time resolution). Enlarged view of one revolution (360°) is shown on the right. Pink, red, and black traces represent raw, median-filtered (current ± 7 frames), and fitted trajectories, respectively. Inset shows the corresponding *x-y* trajectory. Pink lines and red dots represent the raw and median-filtered (current ± 7 frames) coordinates, respectively. We collected 1718 steps from 23 molecules. Other examples of trajectory are shown in Figure S9. (*B*) Distribution of the step size fitted with single Gaussian assuming single peak. The right top values are fitted parameter (peak ± S.D.) and average. The ratio of backward steps was 1.2%. (*C*) Distribution of the duration time before forward step fitted with a single exponential decay function. The right top values are obtained time constant (fitted value ± S.E. of the fit).

**Figure 5.**
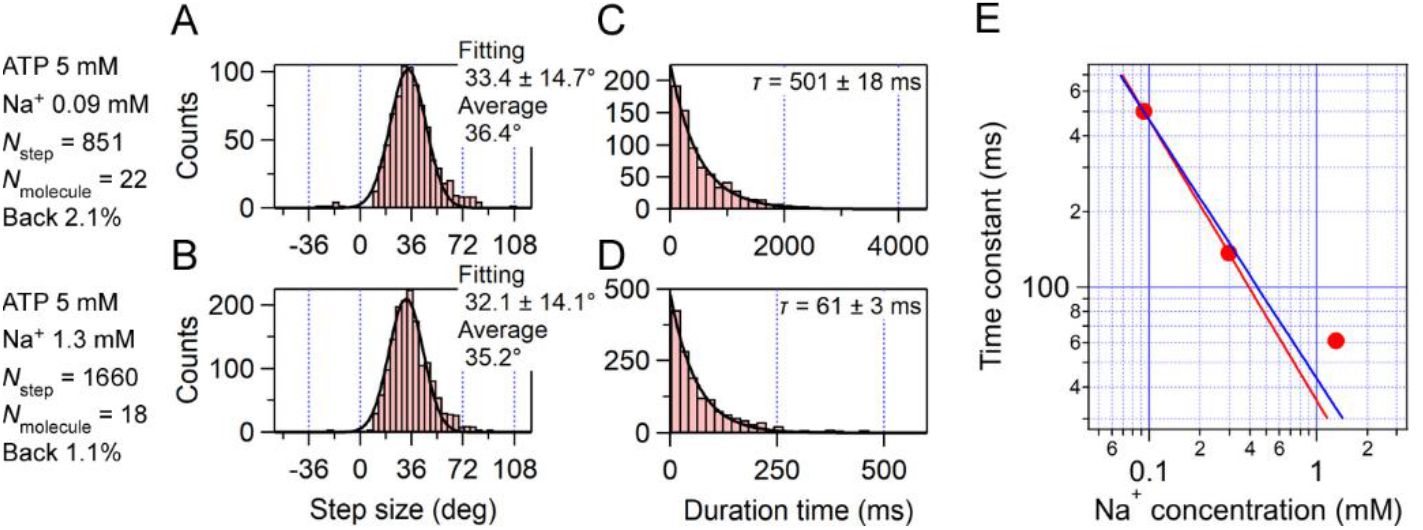
Single-molecule analysis of aE634A at saturated [ATP] (5 mM). Experimental conditions are described on the left of the figures. Examples of trajectory are shown in Figure S10. (*A* and *B*) Distribution of the step size at 0.09 and 1.3 mM Na^+^. Black lines represent fitting with single Gaussians. The right top values are fitted parameters (peak ± S.D.) and averages. (*C* and *D*) Distribution of the duration time before forward step. Black lines represent fitting with single exponential decay functions. The right top values are the obtained time constants (fitted value ± S.E. of the fit). (E) Plot between [Na^+^] (0.09, 0.3, and 1.3 mM) and time constant obtained by the fitting at 5 mM ATP. Solid red line represents a straight line connecting two data points at 0.09 and 0.3 mM Na^+^, and its slope is −1.1. Solid bule line is a result of linear fitting among all three data points. The obtained slope is −1.0.

Next, we observed rotation of aE634A at 0.3 mM Na^+^ and 1 μM ATP with 1,000 fps (Figures 6 and S11). Under this condition, both Na^+^ and ATP bindings are the rate-limiting of rotation (Figures 2B and C). The step-finding algorithm was applied to the median-filtered trace (current ± 7 frames, red line). As a result, the number of detected pauses in a single turn was mainly 13, reflecting 10-pausing positions of EhV_o_ and 3-pausing positions of EhV_1_ (Figure S8C). This result indicates that the pausing positions waiting for Na^+^ and ATP bindings are different in EhV_o_V_1_ complex. Distribution of the step size is shown in Figure 6B. The average value for forward step was 29.9° and distinctly smaller than 36°. When we superimposed two histograms of Figure 4B (0.3 mM Na^+^ and 5 mM ATP) and Figure 6B (0.3 mM Na^+^ and 1 μM ATP) after normalizing the maximum values, difference was obvious (Figure S12). Furthermore, interestingly, in contrast to the condition that Na^+^ binding was the sole rate-limiting factor, the ratio of backward step increased to 6.1%. Therefore, we fitted the distribution with the sum of three Gaussians: one peak in backward (minus) direction and two peaks in forward (plus) direction. Note that one of the forward peaks was fixed at 36°, assuming as the step of EhV_o_. Then, we obtained three peaks at −14.2 ± 6.8, 23.1 ± 10.4, and 36.0 ± 12.8° (peak ± S.D., Figure 6B). For comparison, the fitting with two Gaussians with peaks at minus and plus directions is shown in Figure S13A. Distribution of the duration time before forward step was better fitted with a sum of two exponential decay functions (coefficient of determination *R*^2^ = 0.98, Figure 6C) than with a single exponential decay function (*R*^2^ = 0.94, Figure S13B), and time constants of 135 ± 24 and 642 ± 322 ms (fitted value ± S.E. of the fit) were obtained. These values presumably correspond to the time constants for Na^+^ and ATP bindings, respectively, because the time constant for Na^+^ binding was 137 ms at 0.3 mM Na^+^ and 5 mM ATP (Figure 4C), and that for ATP binding was 476 ms at 1 μM ATP estimated from *k*_on_^ATP^ (2.1 × 10^6^ M^-1^s^-1^, Table 1). These results are consistent with the notion that both Na^+^ and ATP bindings are the rate-limiting of rotation.

**Figure 6.**
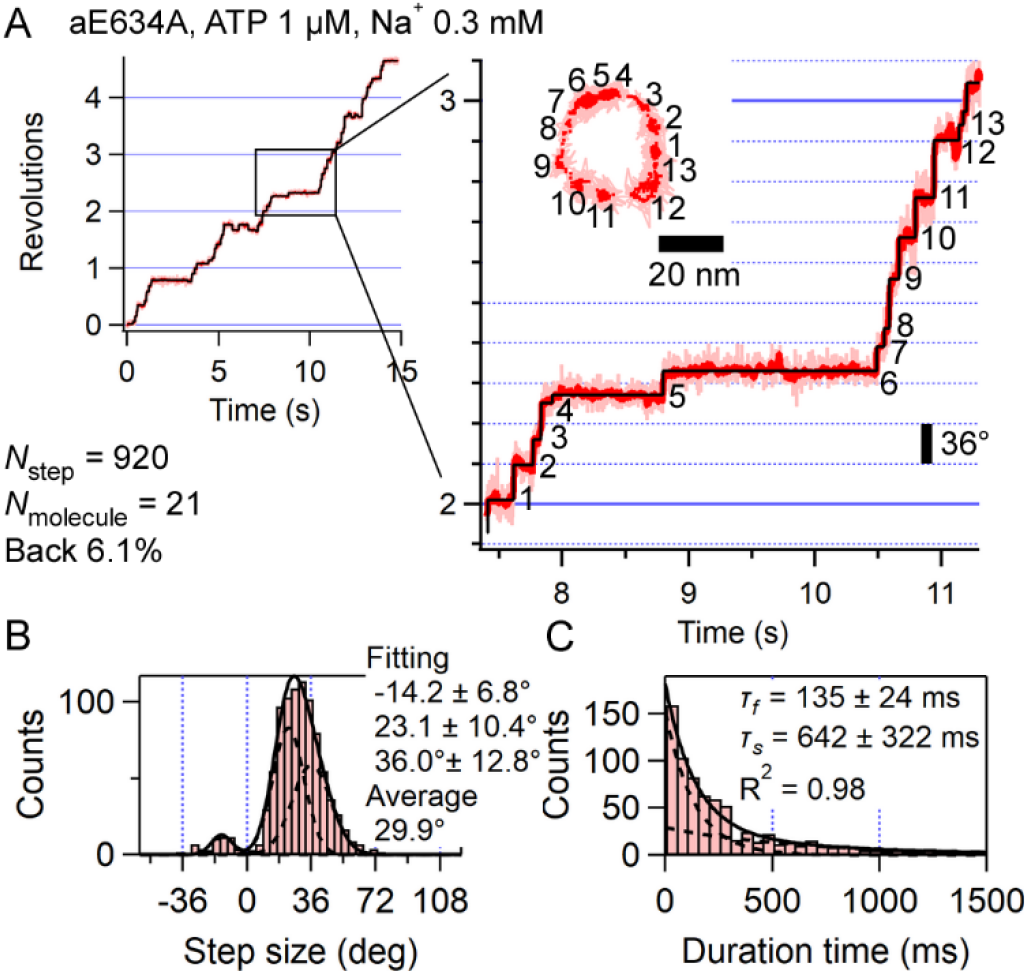
(*A*) Typical trajectory of ATP-driven rotation of aE634A at 1 μM ATP and 0.3 mM Na^+^ recorded with 1,000 fps (1 ms time resolution). Enlarged view of one revolution (360°) is shown on the right. Pink, red, and black traces represent raw, median-filtered (current ± 7 frames), and fitted trajectories, respectively. Inset shows the corresponding *x-y* trajectory. Pink lines and red dots represent the raw and median-filtered (current ± 7 frames) coordinates, respectively. We collected 920 steps from 21 molecules. Other examples of trajectory are shown in Figure S11. (*B*) Distribution of the step size fitted with a sum of three Gaussians: one peak in backward (minus) direction and two peaks in forward (plus) direction, one of which was fixed at 36°, assuming that it was the step of EhV_o_. The ratio of backward steps was 6.1%. (*C*) Distribution of the duration time before forward step fitted with a sum of two exponential decay functions. The right top values are obtained time constants (fitted value ± S.E. of the fit) and the coefficient of determination (*R*^2^) of fitting.

### Backward and recovery steps between adjacent pauses of EhV_o_ and EhV_1_

As described above, frequent backward steps (6.1% of total steps) were observed under the condition that both Na^+^ and ATP bindings are the rate-limiting of rotation (Figure 6B). Typical examples of the backward step are shown in Figure 7. Green, cyan, and purple lines indicate forward step just before the backward step, backward step, and forward step after the backward step (recovery step), respectively. Figures 7B-D show the step size distributions for these events. The average value of backward step size was −18.8° (Figure 7C), which is smaller than the expected step size (36°) of EhV_o_. Because backward steps were frequently observed when not only Na^+^ binding but also ATP binding was the rate-limiting of rotation, it is reasonable to assume that backward steps occur during the pauses waiting for ATP binding. The backward step size smaller than 36° is also consistent with the notion that the pausing positions waiting for Na^+^ and ATP bindings are different in EhV_o_V_1_ complex, and backward steps occur at the intermediate pausing position of EhV_1_ between pausing positions of EhV_o_. The forward steps just before the backward and recovery steps seemed to show two kinds of step sizes, smaller than 36° and close to 36° (Figures 7B and D). The backward and forward steps smaller than 36° suggest the Brownian motion between the adjacent pausing positions waiting for ATP and Na^+^ bindings. Furthermore, 36° forward steps just before the backward steps suggest a standard forward step in EhV_o_ followed by the Brownian backward step to the adjacent pausing position of the EhV_1_, and 36° recovery steps after the backward steps suggest a resumption of the rotation after ATP binding to EhV_1_.

**Figure 7.**
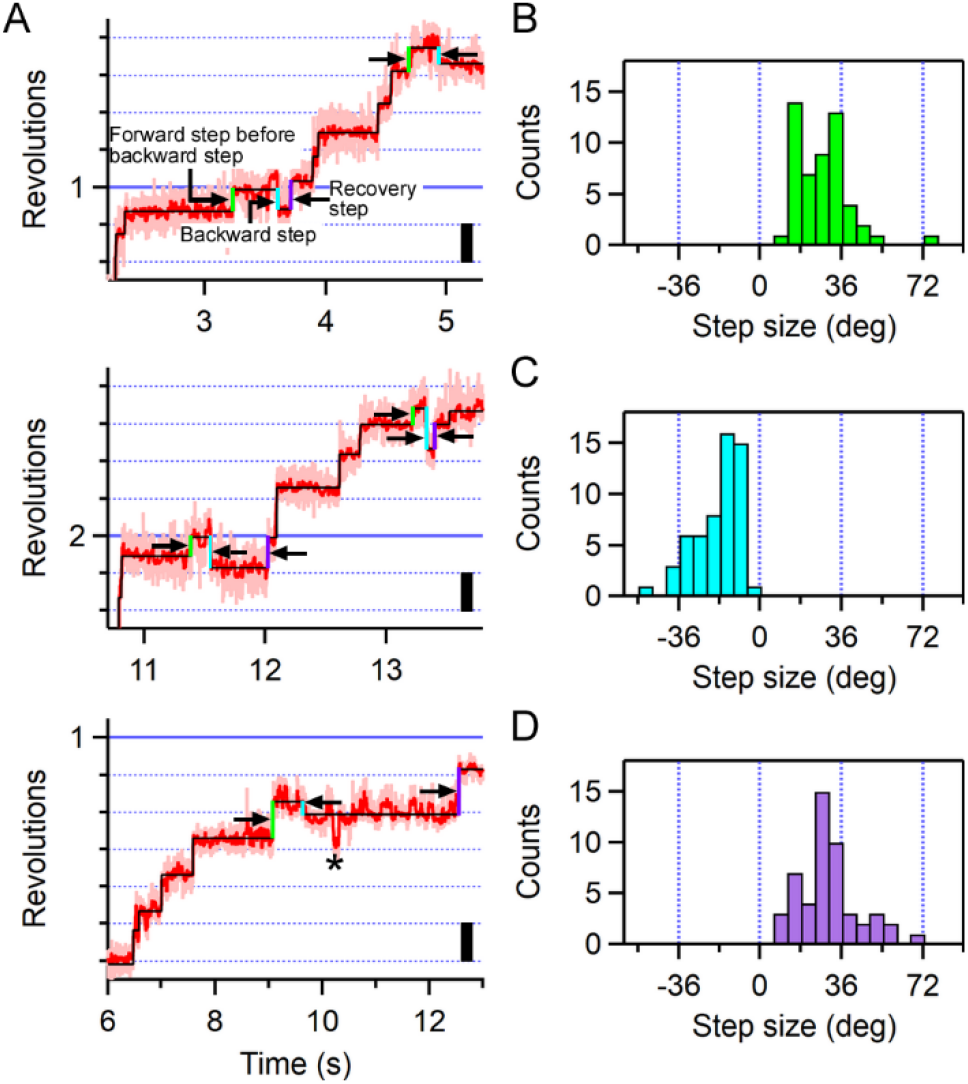
Backward steps of aE634A observed at 1 μM ATP and 0.3 mM Na^+^ with 1,000 fps (1 ms time resolution). (*A*) Examples of trajectories showing the backward steps. The pink, red, and black traces represent the raw, median-filtered (current ± 7 frames), and fitted trajectories of the median-filtered data identified by the algorithm, respectively. The green, cyan, and purple lines indicate forward steps just before backward steps, backward steps, and forward steps just after backward steps (recovery steps), respectively. An asterisk indicates a backward step that is not detected as a step by the algorithm (under-fitting). (*B to D*) Distributions of step size for forward steps just before backward steps, backward steps, and recovery steps, respectively. In *B* and *D*, the distributions seemed to show two peaks at <36° and 36°. In *C*, the peak position was larger than −36° and the average value was −18.8°.

Then, to test the hypothesis whether the backward steps occur equally at three pausing positions of EhV_1_, we prepared other mutants, isolated EhV_1_(BR350K) and EhV_o_V_1_(BR350K) complex. Because an arginine residue in the B-subunit (BArg350) of EhV_1_ plays a crucial role in ATP hydrolysis (Figure S14) (40), the rotation rate of EhV_1_(BR350K) mutant decreases about 100 times than that of wild-type EhV_1_ (44). This property makes ATP cleavage pauses of EhV_1_ much longer in EhV_o_V_1_(BR350K) even at high [ATP], and the pauses of EhV_o_ and EhV_1_ become distinguishable. The rotations of EhV_1_(BR350K) and EhV_o_V_1_(BR350K) observed at 0.3 mM Na^+^ and 5 mM ATP with 1,000 fps are shown in Figure S15. Note that in the isolated EhV_1_(BR350K), His_6_-tag was introduced to the N-terminus of the A-subunit to immobilize the stator A_3_B_3_ ring, and 40-nm AuNP probe was attached to the rotor D-subunit (Figure S14). Three long pauses were observed in both mutants, attributed to ATP cleavage pauses. Similar to our previous study, EhV_1_(BR350K) did not show clear backward steps during long ATP cleavage pauses under this condition (Figure S15A) (44). On the other hand, EhV_o_V_1_(BR350K) exhibited frequent backward and recovery steps during long ATP cleavage pauses (Figure S15B). Distribution of the step size showed two peaks at −11 and 11° and the sizes were smaller than 36° (Figure S15C), and these small backward and recovery steps occurred equally in all three ATP cleavage pauses of EhV_1_.

## Discussion

### ATP-driven rotation of EhV_o_V_1_ tightly coupled with [Na^+^]

In the present study, we directly visualized ATP-driven rotation of EhV_o_V_1_ rate-limited by ion transport. EhV_o_V_1_ used in this study transports Na^+^, while most of the rotary ATPases transport H^+^. This property of EhV_o_V_1_ has advantages to resolve the stepping rotation rate-limited by ion transport in V_o_. Firstly, H^+^ transport in rotary ATPases is thought to be achieved by a Grotthuss mechanism (14, 65), in which H^+^ transfer is not a rate-limiting process. Indeed, the duration time of transient dwells of F_o_ in *E. coli* F_o_F_1_ did not exhibit pH dependence ranging from pH 5 to 9, although the frequency of pause occurrence was highly affected by pH (14). In addition, changes in pH can largely affect protein stabilities and enzymatic reactions, whereas changes in [Na^+^] would mildly affect them. Here, we resolved the pauses and steps of EhV_o_V_1_ by using a mutant in which aGlu634 residue in the stator a-subunit of EhV_o_ was substituted with alanine (aE634A) and found that the duration time before forward steps is inversely proportional to [Na^+^] (Figure 5E). We have previously confirmed that the coupling in EhV_o_V_1_ is retained by using DCCD (*N, N*’-dicyclohexylcarbodiimide) modification assay (16). [Na^+^] dependences of the rotation rate in both wild-type and aE634A (Figure 2C) are also strong evidence for the intact coupling. To the best of our knowledge, the present study first demonstrated ATP-driven rotation of a rotary ATPase rate-limited by the ion transport and tight coupling between ion binding and rotational step, although single-molecule studies of ATP-driven rotation of H^+^-transporting rotary ATPases have been reported (12-14, 66-69).

We found that both wild-type and aE634A mainly shows 10 pauses per single turn and 36° steps (Figures 3 and 4), which reflect the structural symmetry of c_10_-ring under the condition that only Na^+^-binding is the rate-limiting of rotation (0.3 mM Na^+^ and 5 mM ATP). In addition, number of pauses larger or smaller than 10 per single turn was also observed (Figure S8). From the time constants for elementary steps of ATP hydrolysis reaction in EhV_1_ and the time resolution of measurement (0.33 and 1 ms for wild-type and aE634A, respectively), we can consider the case where more than 10 pauses are detected (Figure S8). From the *k*_on_^ATP^ of wild-type and aE634A (2.0×10^6^ and 2.1×10^6^ M^-1^s^-1^, respectively, Table 1), the time constants for ATP binding are estimated to be about 0.1 ms at 5 mM ATP. In addition, by using the isolated EhV_1_, we previously reported that the time constants for ATP cleavage and phosphate release are smaller than 1 ms, and that for ADP release is 2.5 ms. Furthermore, ADP release occurs at different angle from that of ATP binding, while ATP cleavage and phosphate release occur at same angle to ATP binding (44). Considering stochastic nature of pause duration and current time resolution, these elementary events, especially ADP release, can be also detected. If the time resolution and the localization precision is further improved, detection of 16 pauses (10, 3, and 3 pauses for Na^+^ binding, ATP binding, and ADP release, respectively) would become possible, and give us further insight into the energy transduction mechanism of EhV_o_V_1_ complex. Similarly, number of pauses less than 10 can also occur stochastically due to Na^+^ binding events with shorter dwell time than the time resolution. Another possible explanation is stochastic H^+^ transport instead of Na^+^ in EhV_o_V_1_ at low [Na^+^]. It has been reported that H^+^ can bind to the ion-binding site of EhV_o_ c-subunit when [Na^+^] is sufficiently low (42). Because H^+^ binding will not be the rate-limiting factor as described above, the pauses could not be detected at the current time resolution (13). In the present study, we could not distinguish Na^+^ and H^+^ bindings, and assumed that steps are triggered by Na^+^ bindings. Whether or not EhV_o_V_1_ really transports H^+^ under the low [Na^+^] conditions will be discussed elsewhere. Furthermore, we need to mention that the step-finding algorithm we used cannot perfectly eliminate the over-and under-fittings (62, 64).

The *V*_max_^ATP^ of wild-type EhV_o_V_1_ (40.0 ± 0.3 rps) at 300 mM Na^+^ was smaller than that of isolated EhV_1_ (117 ± 3 rps) (Figure 2B and Table 1) (44). The difference is presumably caused by elementary steps of Na^+^ transport in EhV_o_, especially by Na^+^ release, and/or the interaction (molecular friction) between rotor c_10_-ring and stator a-subunit. If we assume that Na^+^ release is the rate-limiting, we can estimate the time constant for Na^+^ release to be 1.6 ms from the difference of the *V*_max_^ATP^ between EhV_o_V_1_ and EhV_1_. Note that in the present study, both entry and exit half-channels of the a-subunit were exposed to the solution with 300 mM Na^+^ because we used detergent-solubilized EhV_o_V_1_. Effect of Na^+^ rebinding from the exit half-channel on the rotation rate of EhV_o_V_1_ is an interesting question that needs to be clarified in future. On the other hand, the *V*_max_^ATP^ of aE634A (4.58 ± 0.04 rps) at 300 mM Na^+^ was approximately one-tenth that of wild-type (40.0 ± 0.3 rps) (Figure 2B and Table 1). Michaelis-Menten kinetics analysis showed that the *k*_on_^ATP^ of aE634A was similar to that of wild-type, although the *k*_on_^Na^ of aE634A significantly decreased compared with that of wild-type (Tables 1 and 2). These results indicate that aE634A mutation has little effect on ATP binding to EhV_1_. The mutated glutamate residue is located on the surface of entry half-channel of a-subunit (Figure 1C), and the decrease in *k*_on_^Na^ is likely caused by the loss of negative charge. Interestingly, this glutamate residue is completely conserved among Na^+^-and H^+^-transporting V-ATPases (Figures S1 and S2) (23, 25, 60, 70). Therefore, it may have a common role in efficient ion uptake for both Na^+^ and H^+^. Consistent with this notion, ion selectivity seems to be determined by the ion-binding site of c-ring rather than the half-channels in a-subunit (71, 72).

The rotation rate of both wild-type and aE634A showed a biphasic response to [Na^+^] (Figure 2C and Table 2). This biphasic response was also reported previously on the ATPase activity of detergent-solubilized EhV_o_V_1_ (16, 61). The Michaelis-Menten parameters obtained in this study are slightly different from those in previous studies, presumably due to differences in experimental conditions, especially pH and [Na^+^] contaminated in the observation buffer. Regarding the Na^+^ contamination in the experimental system, it should be noted that it is difficult to discuss the affinity at the sub-micromolar range since the observation buffer contains at least 90 μM of Na^+^ (Figure S6). The biphasic response to [Na^+^] strongly suggests two different Na^+^-binding sites of EhV_o_V_1_ with largely different affinities. One possible explanation is that these sites are related to the entry and exit half-channels of a-subunit. The high-affinity binding site may correspond to the Na^+^ binding to the c-ring from the entry site because the dissociation constant of Na^+^ to c-subunit in EhV_o_V_1_ complex was reported as 12 μM in the absence of ATP (42). On the other hand, the low-affinity Na^+^ binding site corresponding to *K*_m2_^Na^ has not been identified yet. We assume that the low-affinity site would be physiologically important because an intracellular [Na^+^] in *Enterococcus hirae* has been reported to be several tens of millimolar (36, 39, 73). To clarify the mechanism of the biphasic response of EhV_o_V_1_ to [Na^+^], it will be required to embed EhV_o_V_1_ in the lipid membrane and change the [Na^+^] in the entry and exit sides independently (74). Alternatively, mutagenesis in the Na^+^ exit half-channel of a-subunit will provide new insights into the biphasic response. Recently, Yanagisawa et al. investigated *E. coli* F_o_F_1_ mutants by replacing charged or polar residues in entry and exit half-channels with non-polar leucine residues and successfully revealed that these residues possess optimal p*K*_a_ values for unidirectional H^+^ transfer (14). A similar approach would be also helpful for EhV_o_V_1_.

### Rigid coupling between EhV_o_ and EhV_1_ in the EhV_o_V_1_ complex

By using single-molecule imaging of aE634A under the condition that both Na^+^ and ATP bindings are the rate-limiting of rotation, we visualized the pauses of EhV_o_ and EhV_1_ simultaneously (Figure 6A and S11). We found that aE634A mainly shows 13 pauses per single turn under this condition (Figure S8C). Furthermore, frequent backward and recovery steps smaller than 36° were observed (Figure 7), not only in aE634A but also during three long ATP cleavage pauses of EhV_o_V_1_(BR350K) (Figure S15). These results indicate that pausing positions waiting for Na^+^ and ATP bindings are different, and transitions occur between adjacent pausing positions of EhV_o_ and EhV_1_. These results also indicate that ATP hydrolysis reaction in EhV_1_ is tightly coupled with Na^+^ transport in EhV_o_, and this coupling is not elastic but rigid. Considering our observation system where the stator aA_3_B_3_E_2_G_2_ complex rotates against the rotor dc_10_DF complex immobilized on the glass surface, if the complex shows a fully elastic coupling, the pausing positions of EhV_o_ and EhV_1_ would be superimposed by deformation of the peripheral E_2_G_2_ stalks or the central DF shaft. In this case, single turn (360° revolution) would be divided into two 108° (36°×3) and one 144° (36°×4) with extensive deformation. However, our results clearly showed 10 pauses per single turn (36° step) at high [ATP] and 13 pauses at low [ATP], indicating the rigid coupling in which 120° steps in EhV_1_ are retained in the presence of structural symmetry mismatch between EhV_o_ and EhV_1_.

The different pausing positions between V_o_ and V_1_ have also been reported by a single-molecule study of detergent-solubilized H^+^-transporting *Thermus thermophilus* V/A-ATPase (here termed as TtV_o_V_1_), which acts as ATP synthase (19). During rotation in the ATP-hydrolysis direction, TtV_o_V_1_ showed 30° steps derived from c_12_-ring of TtV_o_ and extra pauses between two adjacent pauses of TtV_o_. These extra pauses were attributed to catalytic dwells of TtV_1_. Furthermore, corresponding structures were also recently revealed by cryo-EM study of nanodisc-embedded TtV_o_V_1_ (23). Zhou et al. resolved structures of two substates in addition to three main states, which reflect three-fold structural symmetry of TtV_1_. Those substates, 1L and 1R, which differ in the relative position of TtV_o_ and TtV_1_, exhibited slight twist of central shaft and deformation of peripheral stalks against state 1. Although it remains uncertain which pauses found in the single-molecule study correspond to those substates, it is intriguing that the angles of TtV_o_ and TtV_1_ do not coincide despite that TtV_o_ has c_12_-ring and no symmetry mismatch with TtV_1_.

The rigid coupling between EhV_o_ and EhV_1_ is distinct from *E. coli, Bacillus* PS3, and yeast mitochondrial F-ATPases, in which cryo-EM studies have proposed the elastic coupling (24, 28, 34). These F-ATPases have c_10_-rings (33, 75) and same structural symmetry mismatches as EhV_o_V_1_. Structural analysis of *E. coli* or *Bacillus* PS3 F_o_F_1_ revealed that three rotational steps were classified into two 108° steps and one 144° step. This significant mismatch was tolerated by the flexible peripheral stalk rather than the central shaft (24, 28, 76). Actually, very recent cryo-EM study of yeast mitochondrial F_o_F_1_ revealed extensive deformation of the peripheral stalk during catalysis (34). In rotary ATPases, number of the peripheral stalks varies from one to three, depending on enzymes (6, 77, 78). *E. coli, Bacillus* PS3 and yeast F_o_F_1_ possess only one peripheral stalk (24, 28), whereas EhV_o_V_1_ has two peripheral stalks which can strongly connect two motor domains (Figure 1A) (25). As a result, EhV_o_V_1_ would rotate rigidly without large and elastic deformations. Consistent with this notion, a cryo-EM study of yeast V_o_V_1_ which has three peripheral stalks showed almost uniform three 120° steps despite having a c_10_-ring (26).

In addition to the number of peripheral stalks, the number of c-subunits in a rotor ring also varies widely from 8 to 17 among the species (56, 57). This makes the issue more complicated, because different number of c-subunits will change the step size and the ion-to-ATP ratio. Furthermore, number of transmembrane helices in a c-subunit monomer also differs among species. For example, although both EhV_o_V_1_ and *E. coli* F_o_F_1_ have c_10_-ring, each c-subunit consists of tetra-and double-transmembrane helices, respectively (24, 53). This results in largely different diameters of c_10_-ring, 8 and 5 nm for EhV_o_V_1_ and *E. coli* F_o_F_1_, respectively (25). Obviously, larger rings would require larger deformations to adjust relative angles between V_o_/F_o_ and V_1_/F_1_ if the elastic coupling is assumed. Another interesting feature of EhV_o_V_1_ with large c-ring is “off-axis” rotation which may affect the coupling between EhV_o_ and EhV_1_ (25). However, the off-axis rotation was not resolved in the present single-molecule experiments. To understand the common and diverse mechanisms of energy transduction in the rotary ATPases, a comprehensive study of various ATPases with rotor c-rings of different c-subunit numbers and sizes will be required.

### Brownian ratchet rotation of EhV_o_

Figure 8 shows the schematic model of the stepping rotation of EhV_o_V_1_ at high [ATP] and low [Na^+^] (Figure 8A), and at low [ATP] and low [Na^+^] (Figure 8B). In the present study, backward steps were rarely observed when [ATP] is high (5 mM) in both wild-type and aE634A (Figures 3B, 4B, 5A, and 5B). In EhV_o_V_1_, the torque for ATP-driven rotation is generated by ATP binding and ADP release in EhV_1_ (44), and the value has been estimated to be 20 pNnm (16). When [ATP] is high, time constants for ATP binding and ADP release are both small, and the torque from EhV_1_ will be applied to EhV_o_ almost constantly. Therefore, EhV_o_V_1_ will show unidirectional rotation without backward steps (Figure 8A). On the other hand, when [ATP] is low (1 μM), where the time constant for ATP binding is large (476 or 642 ms, Table 1 and Figure 6C), frequent backward steps were observed (Figure 7A). The size of the backward steps was much smaller than 36° (14.2 or 18.8°, Figures 6B and 7C). Because no torque is applied when EhV_o_V_1_ is waiting for ATP binding at the pausing position of EhV_1_ (Figure 8B, dark green), it can move to the adjacent backward (and also forward) pausing position of EhV_o_ by the Brownian motion (Figure 8B, blue arrows). Backward and recovery steps smaller than 36° were more frequently observed in EhV_o_V_1_(BR350K), in which ATP cleavage pauses are prominently longer than wild-type (Figure S15). Importantly, backward steps occurred equally during three ATP cleavage pauses of EhV_1_, consistent with the notion that backward steps are driven by thermal fluctuation, not by elastic strain accumulation. These results support Brownian ratchet model of EhV_o_ rotation in addition to the rigid coupling of EhV_o_V_1_. Note that the size of backward steps occurring at three pausing positions of EhV_1_ will not be uniform in the model shown in Figure 8B. However, it was difficult for us to resolve the difference under the current experimental set up.

**Figure 8.**
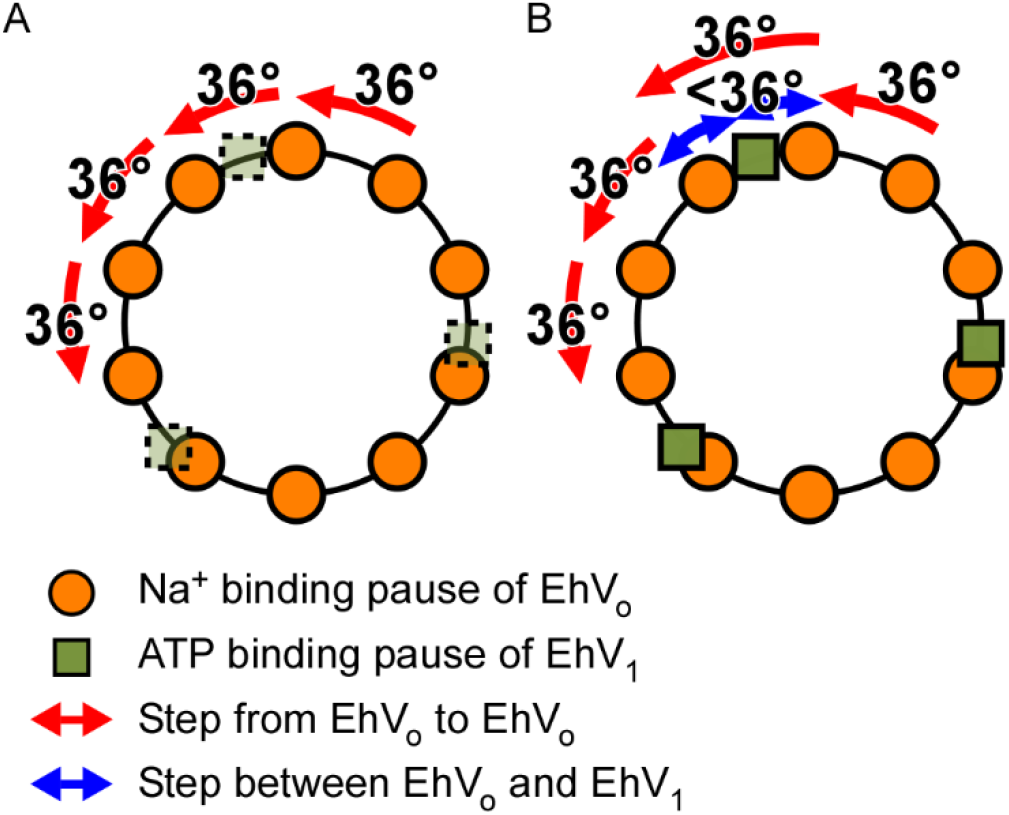
Schematic models of the stepping rotation and rigid coupling of EhV_o_V_1_. The orange circle and dark green square indicate the pausing positions waiting for Na^+^ binding to the EhV_o_ and ATP binding to the EhV_1_, respectively. The red arrows indicate the 36º steps between adjacent pausing positions for the EhV_o_. The blue arrows indicate the backward and forward steps smaller than 36º between adjacent pausing positions for the EhV_o_ and EhV_1_. (*A*) The condition that only Na^+^ binding to the EhV_o_ is the rate-limiting factor. In this condition, the pauses waiting for ATP binding to the EhV_1_ are too short to be detected, and the EhV_o_V_1_ rotates unidirectionally without backward steps. (*B*) The condition that both Na^+^ and ATP bindings are the rate-limiting factors. The pausing positions waiting for ATP binding are visualized and then 13-pausing positions are detected per single turn. Because no torque is generated during the pauses waiting for ATP binding to the EhV_1_, the EhV_o_V_1_ rotates to the backward and forward pausing positions of the EhV_o_ driven by the Brownian motion. As a result, backward and forward steps smaller than 36º are observed.

In our previous study of the isolated EhV_1_, no backward steps are observed in ATP-driven rotation except under extreme experimental conditions (44). Therefore, backward and forward steps smaller than 36° observed in EhV_o_V_1_ are caused by EhV_o_, and are presumably coupled with the Na^+^ binding/release to/from c-subunit through the two half-channels of a-subunit. A recent single-molecule study observed similar small backward steps of 11° in *E. coli* F_o_F_1_ during ATP-driven rotation and successfully revealed that backward steps are related to H^+^ translocation between c-ring and half-channels of a-subunit (14). In the study of *E. coli* F_o_F_1_, small backward step was attributed to the fact that *E. coli* F_o_F_1_ functions as ATP synthase. In the case of EhV_o_V_1_, because backward steps of 36° or larger than 36° rarely occurred, backward flow of Na^+^ seems to be suppressed. The detailed mechanisms of the backward step may be different by ATP synthesis capacity, size of the c-ring, and ion species being transported. In the present study, however, the electrochemical potential of Na^+^ could not be applied because detergent-solubilized EhV_o_V_1_ was used. To address the question of whether the electrochemical potential of Na^+^ can drive rotation of EhV_o_V_1_ in opposite direction and ATP synthesis, our next targets are single-molecule imaging and biochemical assay of EhV_o_V_1_ embedded in the lipid membrane.

## Materials and Methods

### Sample preparation

The construction of the expression plasmid for wild-type EhV_o_V_1_ (pTR19-EhV_o_V_1_) was reported previously (16). For the plasmid construction of aE634A, EhV_o_V_1_(BR350K), and EhV_1_(BR350K), PCR-based site-directed mutagenesis was performed by KOD one PCR Master Mix (Toyobo) with pTR19-EhV_o_V_1_ as a template. For expression and purification of wild-type, aE634A, EhV_o_V_1_(BR350K), and EhV_1_(BR350K), the procedures described in our previous study were used with some modifications (16, 44). Details are described in the SI Appendix.

### Single-molecule imaging

Gold nanoparticle (AuNP) with a diameter of 40 nm (EMGC40, BBI) was biotinylated with biotin-alkane-thiol (HS-C11-EG3, Surfmods) and coated by Streptavidin (PRO791, PROSPEC) as described in the previous reports (44, 79). We followed the setting of a total internal reflection dark-field microscope system based on a high-speed CMOS camera and an inverted microscope as described in our previous studies with some modifications (44). Details are described in the SI Appendix.

### ICP-OES measurement

To estimate the concentrations of contaminated Na^+^ in the observation buffers, we measured emission spectra using an inductivity coupled plasma optical emission spectrometer (ICP-OES; 5100 ICP-OES, Agilent Technologies). The emission signal at λ = 589.592 nm was used for analysis. The flow rates of plasma and assist gas (Ar) were 14 and 1.2 L/min, respectively. The standard addition method was used to avoid physical and ionization interferences. Namely, sodium standard solution (FUJIFILM Wako) was added to the observation buffers in the range of 0.1 to 10 mg/L (4.3 to 430 μM) as a final concentration of Na^+^. The calibration curve was extrapolated and the absolute value of the X-intercept was taken as the concentration of contaminated Na^+^.

## Acknowledgments

We thank Monique Honsa for the advice about data analysis, Kazuyoshi Murata, Chihong Song, and Raymond N. Burton-Smith for providing us the structural data and their fruitful discussion, Yayoi Kon for her grateful technical support, and all laboratory members for helpful discussion and technical advice. This work was supported by the Grant-in-Aid for Scientific Research on Innovative Areas “Molecular Engine” (JP19H05380 to H.U., JP18H05425 to T.M., and JP18H05424 to R.I.), the Grants-in-Aid for Scientific Research (JP21H02454 to R.I., and JP21K15060 and JP20J01316 to A.O.) from the Ministry of Education, Culture, Sports, Science, and Technology of Japan, and by the NINS program for cross-disciplinary study (Grant Number 01312001 to A.O.). A part of this work was performed with the aid of Instrument Center of the Institute for Molecular Science.

