## Supplementary Information for "Direct Observation of Stepping Rotation of V-ATPase Reveals Rigid Coupling between V_o_ and V_1_ Motors"

*Ryota Iino

**This PDF file includes:**

Supplementary text
Figures S1 to S15 (not allowed for Brief Reports)
SI References

**Supplementary Information Text**

### Materials and Methods

*Sample preparation*

For expression and purification of the wild-type, $EhV_oV_1$(aE634A), and $EhV_oV_1$(BR350K) the procedures described in our previous study(1, 2) were used with some modifications. Namely, proteins were expressed in *E. coli* strain C41(DE3) cells. Cells were cultivated in Super Broth medium (32 g/L tryptone, 20 g/L yeast extract, and 5 g/L NaCl) containing 100 mg/L ampicillin, and expression was induced with 1.0 mM isopropyl-β-thiogalactopyranoside at 37°C for 20 hours. The cells were collected and suspended in buffer A (10 mM HEPES-KOH (pH 7.5), 5 mM $MgCl_2$, 10% glycerol) and then disrupted by ultrasonication. The cell debris were removed by centrifugation (21,000×g, 30 min at 4°C) and the membrane fraction was collected by ultracentrifugation (100,000×g, 60 min at 4°C). The precipitate was suspended in buffer B (50 mM potassium phosphate (pH7.5), 100 mM KCl, 5 mM $MgCl_2$, 20 mM imidazole, and 10% glycerol) by gently pipetting in the ultrasonic bath. The suspension was solubilized with 2% n-dodecyl-β-D-maltoside (DDM) (Sigma) by rotating for 30 min at 4°C. The solubilized fraction was collected by ultracentrifugation (100,000×g, 30 min at 4°C) and the supernatant was loaded on $Ni^{2+}$-NTA agarose (superflow, QIAGEN) column equilibrated with the buffer B containing 0.05% DDM. After column wash extensively with the buffer B containing 0.05% DDM, $EhV_oV_1$ was eluted with buffer C (50 mM potassium phosphate (pH7.5), 50 mM KCl, 5 mM $MgCl_2$, 300 mM imidazole, 10% glycerol, and 0.05% DDM) and concentrated to <500 µL by ultrafiltration (Vivaspin6 30,000 MWCO). The concentrated sample was applied onto Superdex200 increase 10/300 GL column equilibrated with buffer D (50 mM Tris-HCl (pH 7.5), 5 mM $MgCl_2$, 10% glycerol, and 0.05% DDM) by using HPLC system (JASCO). The fractions which contain the $EhV_oV_1$ complex were collected and stored at -80°C until use. The concentration of purified $EhV_oV_1$ was estimated by the molar extinction coefficient; $\varepsilon_{280nm}$ = 512,860 $M^{-1}cm^{-1}$.

*Single-molecule imaging and image data analysis*

Flow cell was prepared by greasing a thin spacer on top of the $Ni^{2+}$-NTA coated cover glass (24 mm × 32 mm, Matsunami Glass) and placing a smaller cover glass on top (18 mm × 18 mm, Matsunami Glass). The 8 µL of $EhV_oV_1$ (5 – 10 nM) diluted with Observation buffer A (20 mM KPi or 50 mM Bis-Tris (pH 6.5), 230 mM KCl, and 0.05% DDM) was injected into the flow cell. After 10 min, 40 µL of 5 mg/mL BSA solution dissolved in the Observation buffer A was applied to the flow cell. To avoid the contamination of $Na^+$, BSA (Wako) was desalted beforehand by desalting column (PD-10, GE healthcare). After 5 min, 8 µL of streptavidin-coated AuNPs 40 nm suspension in the Observation buffer A supplemented with 5 mg/mL of BSA were injected into the flow cell. After 10 min, to remove the unbound AuNPs and to start ATP-driven rotation, 40 µL of Observation buffer B (20 mM KPi or 50 mM Bis-Tris (pH 6.5), and 50 mM KCl, 2 mM $MgCl_2$, and ATP-regeneration system (2.5 mM phosphoenolpyruvate (Sigma) and 0.1 mg/mL pyruvate kinase (Sigma) with various concentrations of ATP and NaCl) were injected to the flow cell.

Single-molecule rotation assay was performed with a total internal reflection dark-field microscope system at 25°C (2, 3). This system was constructed by an inverted microscope (IX70, Olympus) and a 60× objective lens (numerical aperture = 1.49, Olympus) with an ultra-stable microscope stage (KS-O, Ikeda-rika) and 532-nm CW laser (LDC-2500S, Photop), which was introduced from the side port of the microscope. The laser light was total internal reflected at the interface between the cover glass and the buffer solution and illuminates the sample fixed on the $Ni^{2+}$-NTA coated glass surface. The scattered light from the AuNP was collected through the same objective lens and the perforated mirror and imaged with a high-speed CMOS camera (Fastcam AX100, Photron) at 1,000 or 3,000 frames per second (fps) with a pixel size of 79.7 nm in both x- and y-axis, and recorded as an 8-bit image sequence file (AVI format). The power density of the laser was 1.6 and 4.5 µW/µm$^2$ at the sample plane for 1,000 and 3,000 fps, respectively. The localization precision in the x- and y-axis were 0.54 and 0.58 nm and 0.63 and 0.64 nm for 1,000 and 3,000 fps, respectively (Figure S2). Image data were analyzed by ImageJ software (NIH). The centroids (*x*- and *y*-coordinates) of the image sequences of single AuNPs were analyzed for each frame and converted to the trajectory of rotation. We collected and analyzed the molecules which continuously rotate for more than 5 seconds (aE634A mutant) or more than 5 revolutions (wild-type) as criteria. For analysis of step size and duration time, the trajectory of rotation was mediansmoothed with current ± 4 (0.3 mM $Na^+$, wild-type), ± 7 (0.3 and 1.3 mM $Na^+$, aE634A), or ± 15 frames (for 0.09 mM $Na^+$, aE634A) and applied a step-finding algorithm developed by Kerssemakers (4).

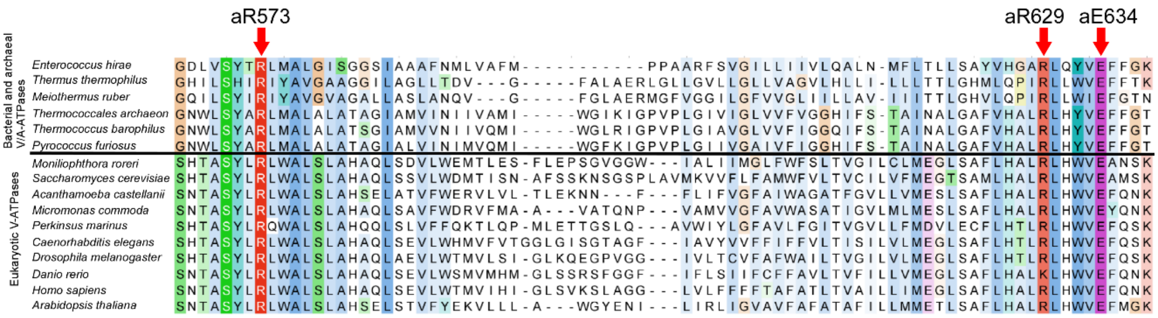


**Fig. S1.** Alignment of amino acid sequences of the a-subunit of V- or A-ATPases from 16 different species created by Clustal Omega (5) with reference to L. Zhou et al., *Science* (2019) (6). The top is the sequence of the a-subunit of *Enterococcus hirae* V-ATPase (amino acid residues from positions 566 to 638). The highly conserved residues are marked with dark colors. The positions of two conserved arginine residues (aR573 and aR629 for *Enterococcus hirae*) located at the interface between two half-channels and a conserved glutamate residue (aE634) located on the surface of the entry half-channel (Figure 1C) are indicated by red arrows.

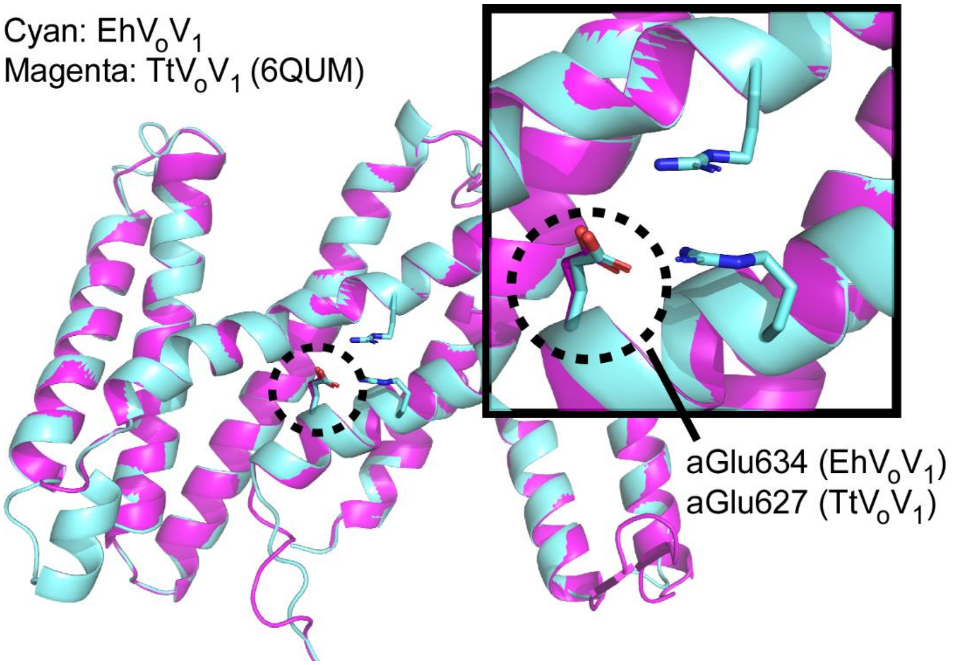


**Fig. S2.** Homology modeling structure of the a-subunit of $EhV_o$ (cyan, amino acid residues from positions 364 to 654) (7) and the structure of the a-subunit of $TtV_o$ used as a template (magenta, amino acid residues from positions 346 to 649, PDB ID: 6QUM) (6). The inset shows an enlarged view, and the dotted circle indicates the conserved glutamate residues.

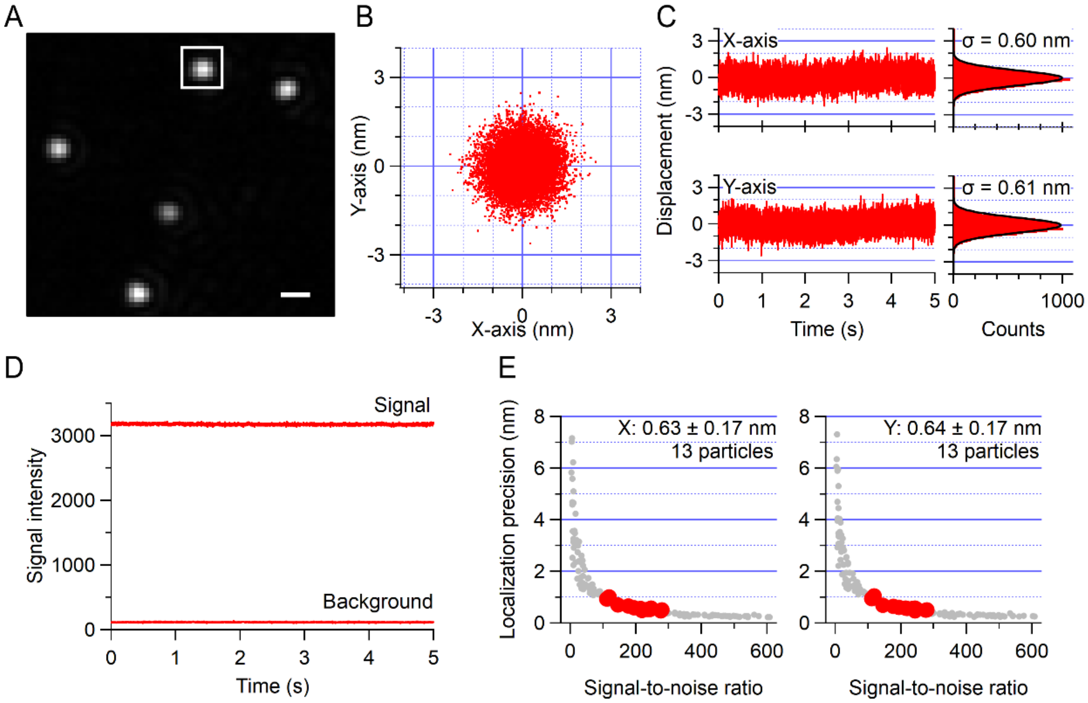


**Fig. S3.** A series of experiment and analysis to determine the localization precision. (*A*) Dark-filed image of 40 nm AuNPs with a pixel size of 79.7 nm/pixel. The scale bar is 500 nm. The scattering image was recorded with 4.5 μW/μm$^2$ and 3,000 fps for 5 seconds. A white square indicates the region of interest (ROI, 10 × 10 pixels) to analyze the centroid coordination and signal intensity of single AuNP. (*B*) Two-dimensional plot for the centroid of a 40 nm AuNP. (C) Time courses of displacement (left) and histograms with Gaussian fit (right) for both X- (top) and Y-axis (bottom). The standard deviation obtained from the Gaussian fitting to the distribution was defined as the localization precision. (*D*) Time courses of signal intensity within the ROI. Background intensity was obtained from the region near the analyzed particle where no particles were present. (*E*) Relationships between localization precision and signal-to-noise ratio for both X- (left) and Y-axis (right). The red and gray circles represent the present results and data reported by Ando et al. (3), respectively.

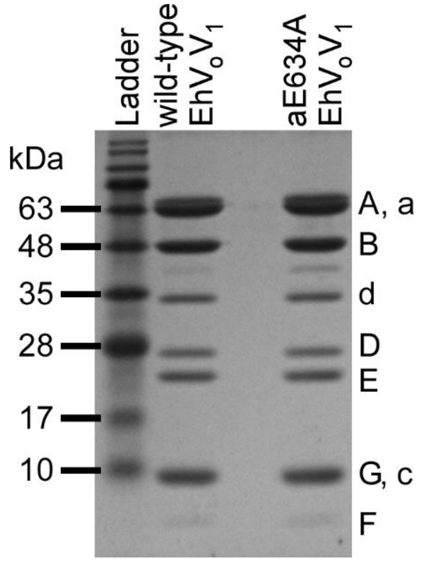


**Fig. S4.** SDS-PAGE of the wild-type and aE634A. 15% gel was used. The band position of each subunit was assigned based on previous reports (1).

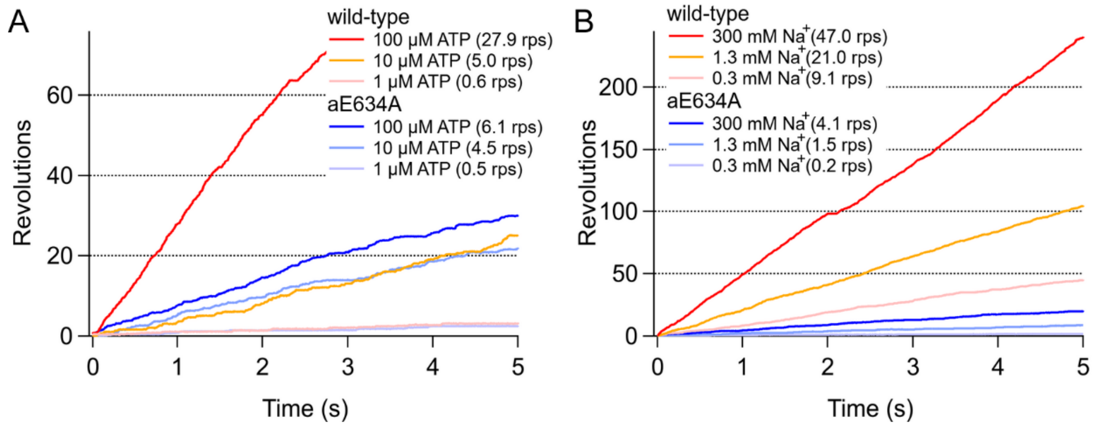


**Fig. S5.** Typical examples of the time course of rotation observed at 300 mM $Na^+$ and different [ATP]s (indicated at the right top of the graph) (*A*) and at 5 mM ATP and different [$Na^+$]s (indicated at the left top of the graph) (*B*). The rotation rate was calculated from the slope of the time course. Data for wild-type and aE634A was recorded at 3,000 and 1,000 fps, respectively.

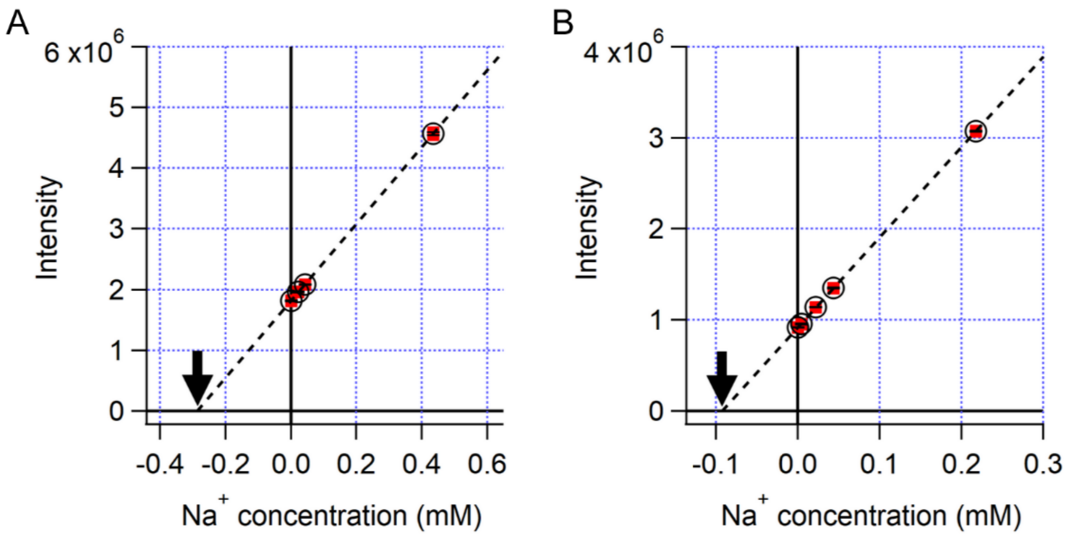


**Fig. S6.** Calibration curve of [$Na^+$] measured by ICP-OES with the standard addition method. We measured two different observation buffers; (*A*) 20 mM potassium phosphate (pH 6.5), 50 mM KCl, 5 mM ATP-$Mg^{2+}$, 2 mM $MgCl_2$, 2.5 mM PEP, 0.1 mg/mL PK, and 0.05% DDM (*B*) 50 mM Bis-Tris (pH6.5), 50 mM KCl, 5 mM ATP-Tris, 7 mM $MgCl_2$, 2.5 mM PEP, 0.1 mg/mL PK, and 0.05% DDM. The red square, black circle, black bar, and dotted line represent the signal intensity at 589.592 nm, the averaged value, the standard deviation (N=3), and fitting line, respectively. The coefficient of determination was 0.999 for both (*A*) and (*B*). The fitting lines are extrapolated and the black arrowheads show the X-intercept. We estimated the amount of contaminated $Na^+$ in each buffer from the absolute value of the X-intercept; 0.3 mM and 0.09 mM for (*A*) and (*B*), respectively.

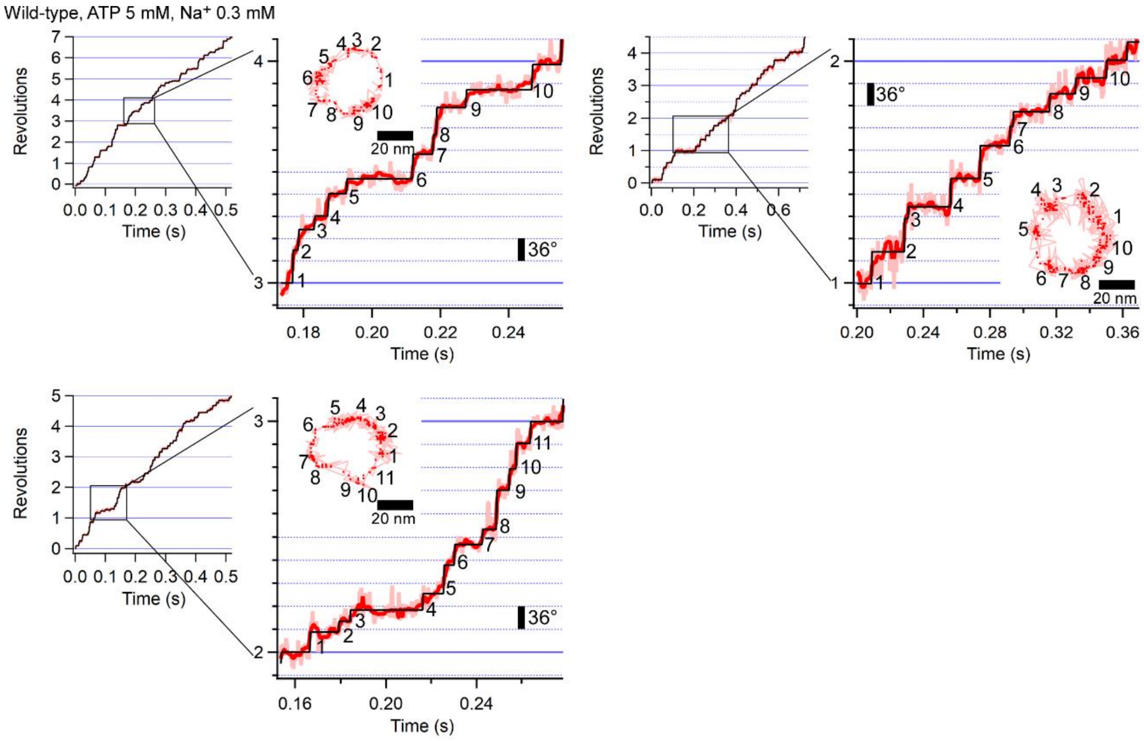


**Fig. S7.** Examples of trajectory of ATP-driven rotation of wild-type at 5 mM ATP and 0.3 mM $Na^+$ recorded with 3,000 fps (0.33 ms time resolution). Examples of three different molecules are shown. The enlarged view of one revolution (360°) is shown on the right. The pink, red, and black traces represent the raw, median-filtered (current ± 4 frames), and fitted trajectory of the median-filtered data identified by the algorithm (4), respectively. The inset shows the corresponding x-y trajectory. The pink lines and red dots represent the raw and median-filtered (current ± 4 frames) coordinates, respectively.

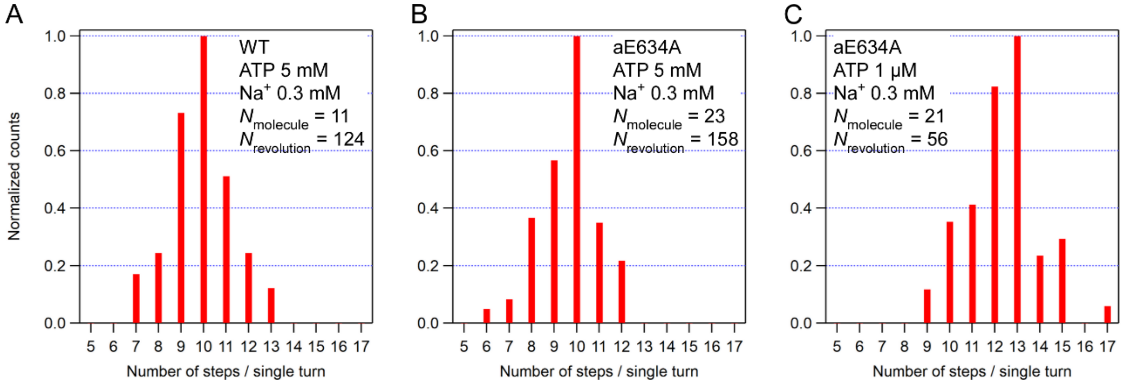


**Fig. S8.** Number of detected pauses per a single turn of (*A*) wild-type at 5 mM ATP and 0.3 mM $Na^+$ corresponding to Figures 3 and S6, (*B*) aE634A at 5 mM ATP and 0.3 mM $Na^+$ corresponding to Figures 4 and S9, and (*C*) aE634A at 1 μM ATP and 0.3 mM $Na^+$ corresponding to Figures 6 and S11.

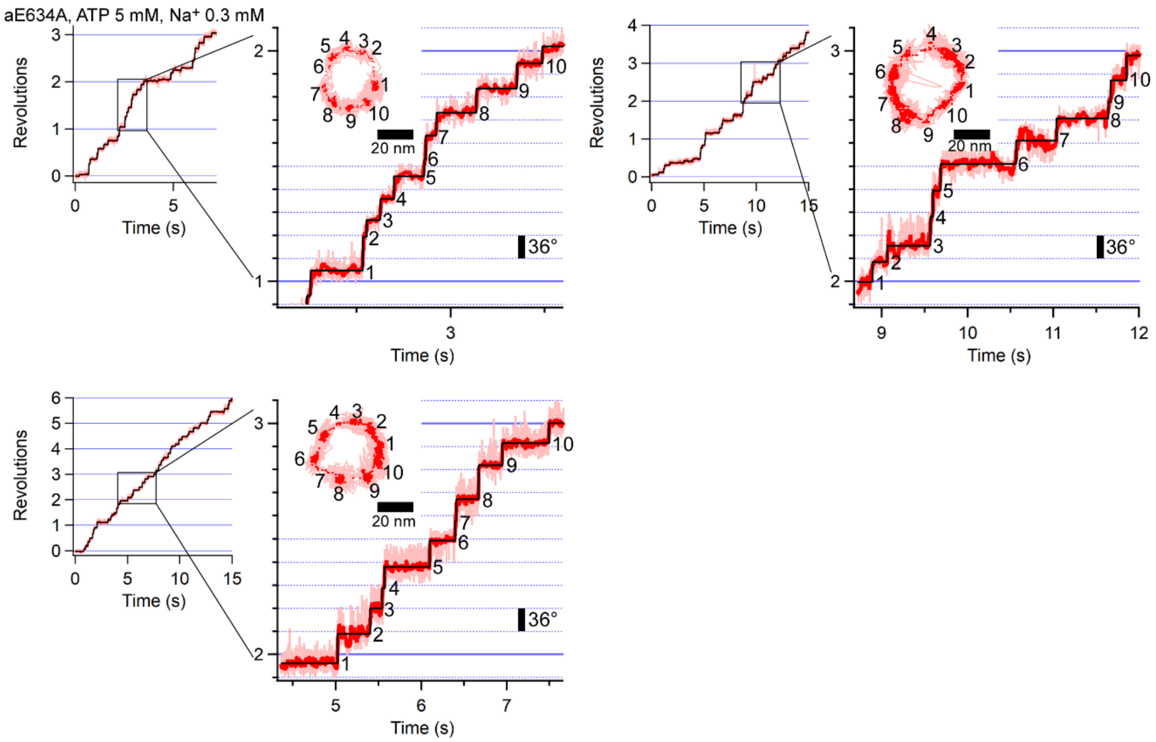


**Fig. S9.** Examples of trajectory of ATP-driven rotation of aE634A at 5 mM ATP and 0.3 mM $Na^+$ recorded with 1,000 fps (1 ms time resolution). Examples of three different molecules are shown. The enlarged view of one revolution (360°) is shown on the right. The pink, red, and black traces represent the raw, median-filtered (current ± 7 frames), and fitted trajectory of the median-filtered data identified by the algorithm (4), respectively. The inset shows the corresponding x-y trajectory. The pink lines and red dots represent the raw and median-filtered (current ± 7 frames) coordinates, respectively.

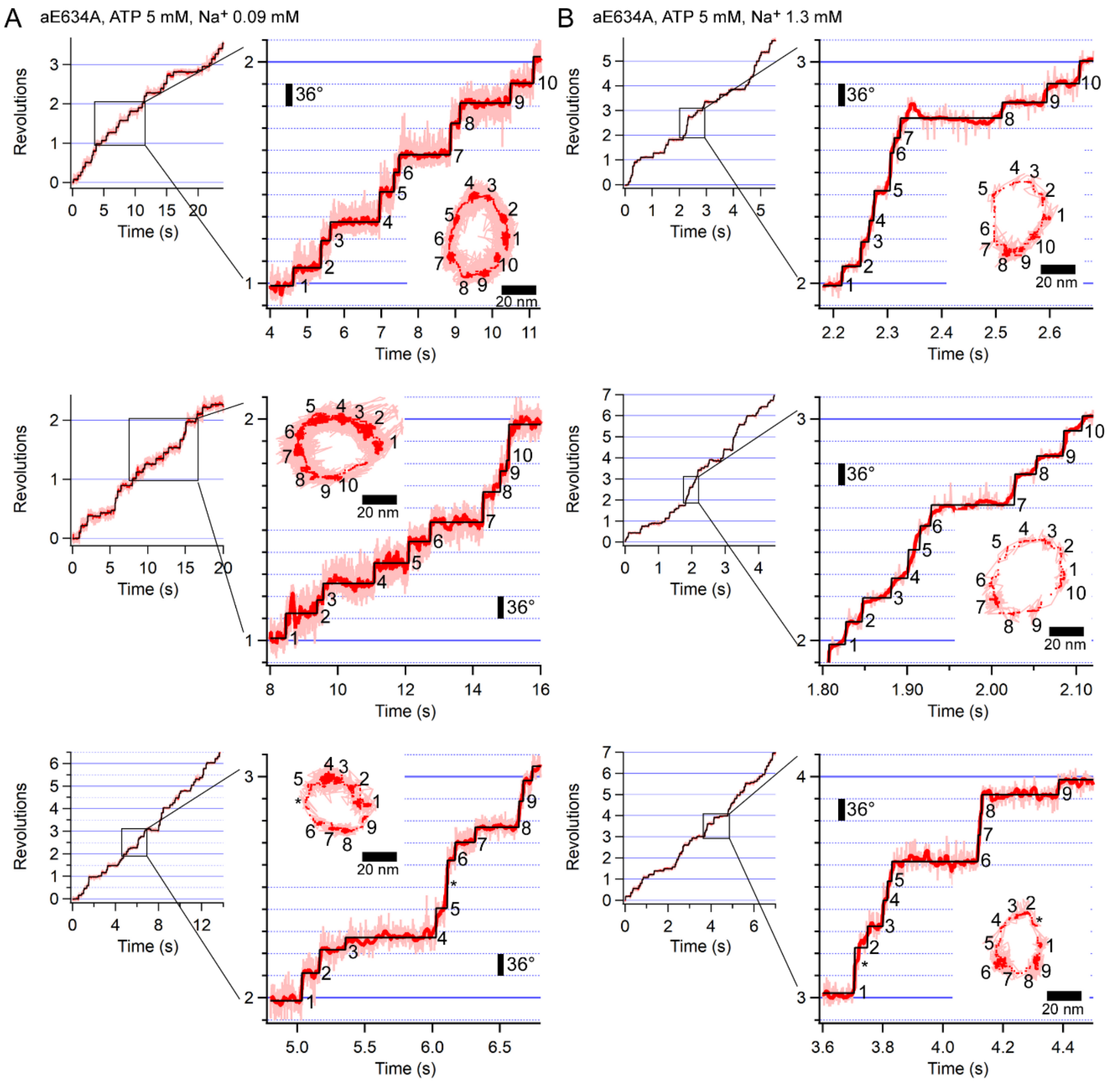


**Fig. S10.** Examples of trajectory of ATP-driven rotation of aE634A at 5 mM ATP and 0.09 mM $Na^+$ (*A*) and 5 mM ATP and 1.3 mM $Na^+$ (*B*) recorded with 1,000 fps (1 ms time resolution). Examples of three different molecules are shown for each experimental condition. The enlarged view of one revolution (360°) is shown on the right. The pink, red, and black traces represent the raw, median-filtered (current ± 15 frames in *A* and ± 7 frames in *B*), and fitted trajectory of the median-filtered data identified by the algorithm (4), respectively. The inset shows the corresponding x-y trajectory. The pink lines and red dots represent the raw and median-filtered (current ± 15 frames in *A* and ± 7 frames in *B*) coordinates, respectively. The short pauses where the algorithm apparently failed to detect are marked with an asterisk.

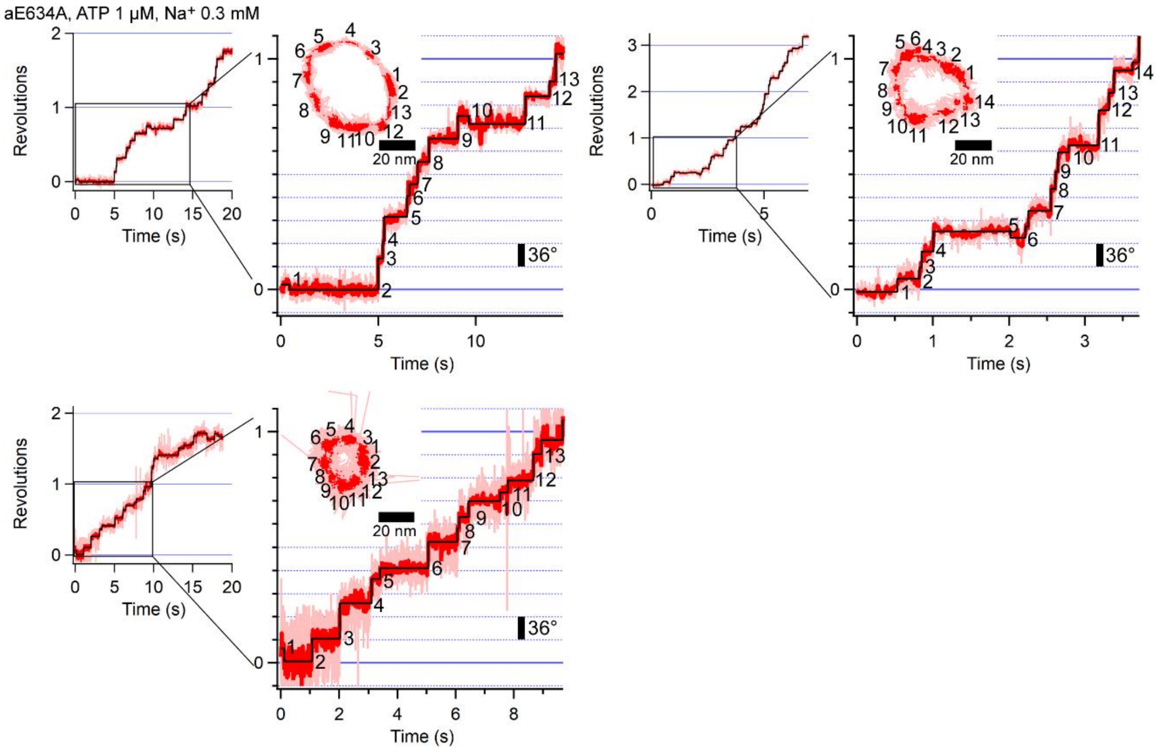


**Fig. S11.** Examples of trajectory of ATP-driven rotation of aE634A at 1 μM ATP and 0.3 mM $Na^+$ recorded with1,000 fps (1 ms time resolution). Examples of three different molecules are shown. The enlarged view of one revolution (360°) is shown on the right. The pink, red, and black traces represent the raw, median-filtered (current ± 7 frames), and fitted trajectory of the median-filtered data identified by the algorithm (4), respectively. The inset shows the corresponding x-y trajectory. The pink lines and red dots represent the raw and median-filtered (current ± 7 frames) coordinates, respectively.

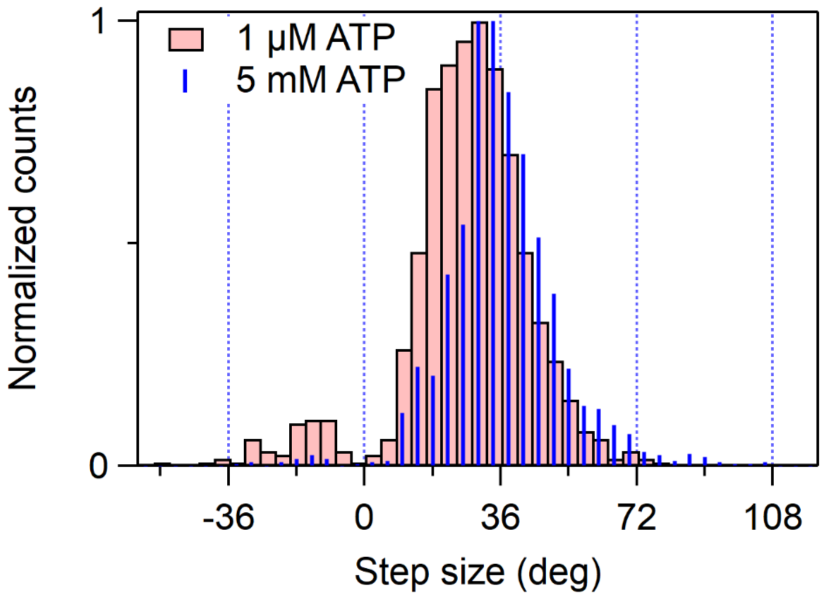


**Fig. S12.** The step size distributions of aE634A at 5 mM ATP and 0.3 mM $Na^+$ (blue) and 1 μM ATP and 0.3 mM $Na^+$ (pink). The histograms for the two conditions were superimposed after the normalization of the maximum values.

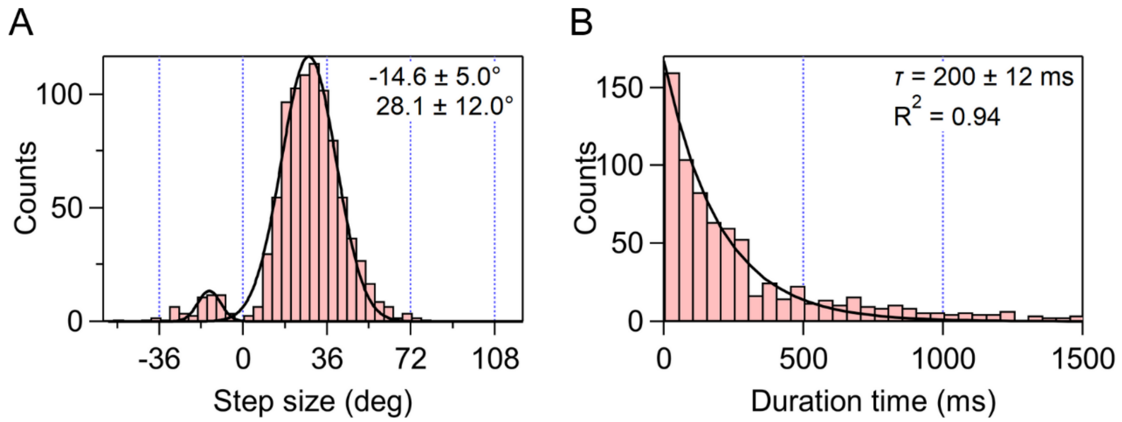


**Fig. S13.** Distributions of the step size (*A*) and the duration time (*B*) before forward steps of aE634A at 1 μM ATP and 0.3 mM $Na^+$. A single component was assumed in the forward steps.

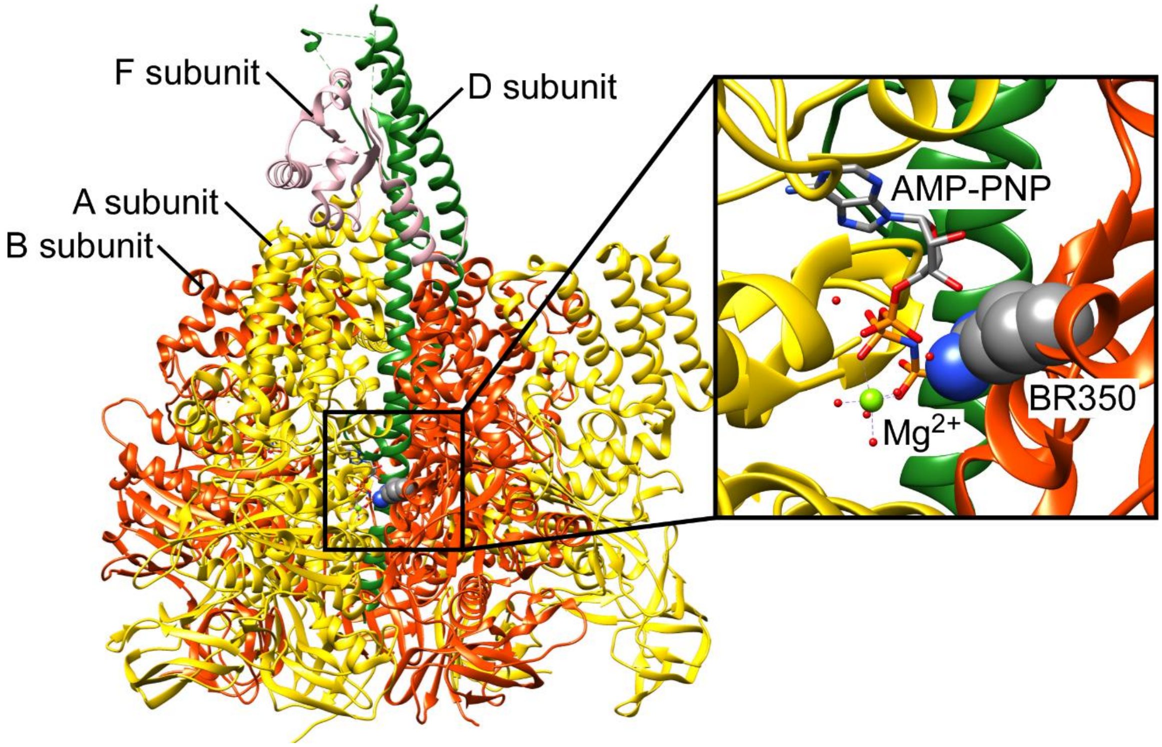


**Fig. S14.** Crystal structure of $EhV_1$ $A_3B_3DF$ complex (PDB ID: 3VR6). The enlarged view is a nucleotide-binding site. In the enlarged view, A and B subunits except for the foreground are shown transparently. The bound AMP-PNP (adenosine 5’-(β,γ-imino)triphosphate), which is an ATP analog, and the Arg350 residue in B-subunit are depicted in stick and sphere format, respectively. Green and red balls are $Mg^{2+}$ and water molecules, respectively.

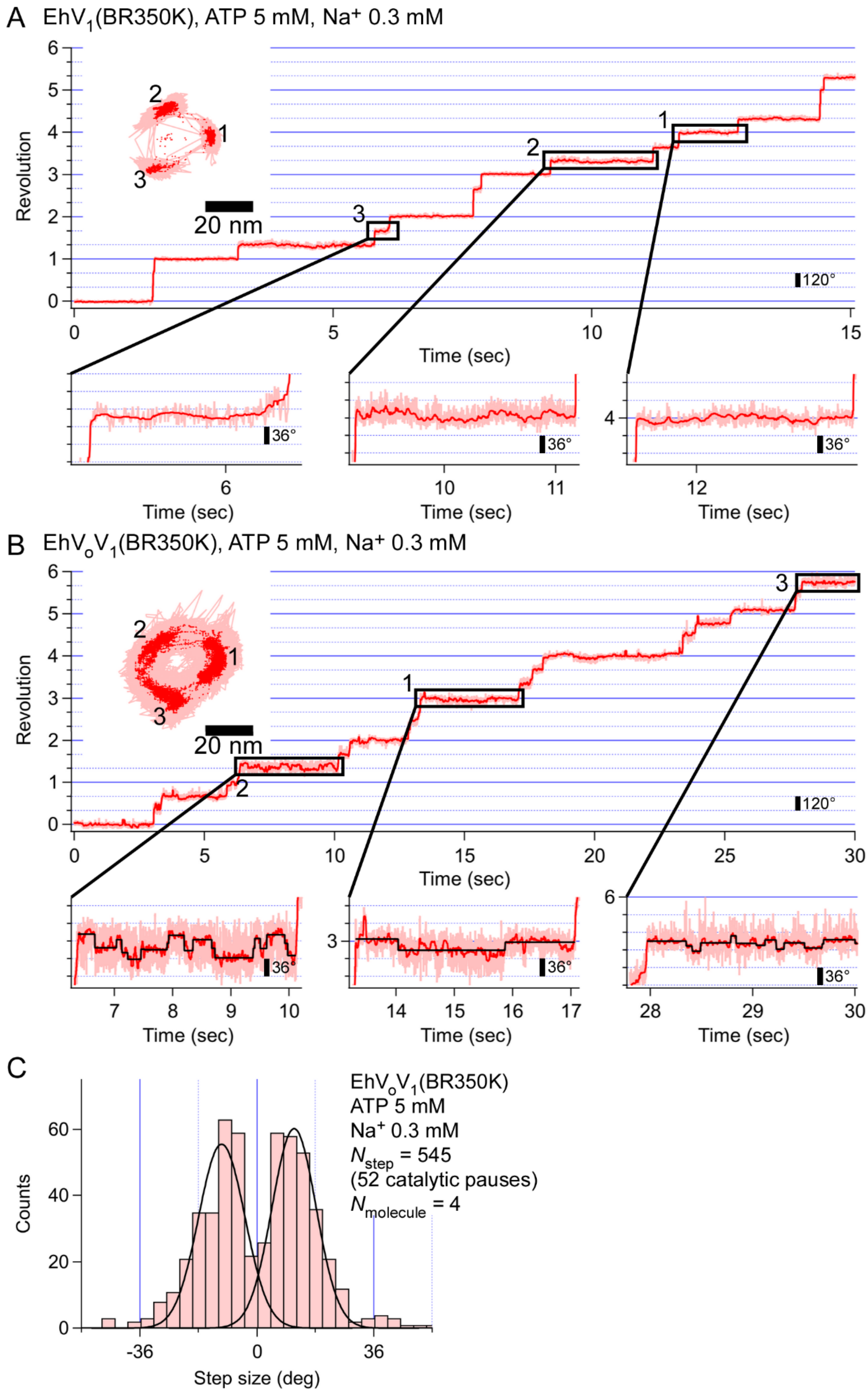

A EhV1(BR350K), ATP 5 mM, Na+ 0.3 mM
Revolution
20 nm
120°
Time (sec)
36°
Time (sec)
B EhVoV1(BR350K), ATP 5 mM, Na+ 0.3 mM
Revolution
20 nm
120°
Time (sec)
36°
Time (sec)
C
EhVoV1(BR350K)
ATP 5 mM
Na+ 0.3 mM
Nstep = 545
(52 catalytic pauses)
Nmolecule = 4
Counts
Step size (deg)

**Fig. S15.** Single-molecule analysis of $EhV_1$(BR350K) (*A*) and $EhV_oV_1$(BR350K) (*B*) at 5 mM ATP and 0.3 mM $Na^+$ recorded at 1,000 fps (1 ms time resolution). An inset on the left top represents the corresponding *x*-*y* trajectory. The pink and red traces represent the raw and median-filtered (current ± 15 frames) data, respectively. The enhanced three pauses are the ATP cleavage pause of $EhV_1$ moiety. The three enlarged views on the bottom show ATP cleavage pauses at three different positions of $EhV_1$ as noted by the numbers. Frequent backward and recovery steps were observed in $EhV_oV_1$(BR350K) and were identified by the step-finding algorithm (4) (black trances in *B*, bottom panels). (*C*) Step size distribution during ATP cleavage pauses of $EhV_oV_1$(BR350K) detected by the algorithm. 52 ATP cleavage pauses and 545 steps from 4 different molecules were analyzed. Data were fitted by two Gaussians. The obtained peak positions were -11 and 11°.
